# The N-terminus of SAS-1 promotes microtubule-dependent centriolar satellite formation

**DOI:** 10.64898/2026.02.27.708307

**Authors:** Asli Beril Tiryakiler, Siti Zakiah Abdul Talib, Andreia Filipa Henriques Soares, Fatma Tutku Altug, Astrid Heim, Esther Zanin, Tamara Mikeladze-Dvali

## Abstract

Centriolar satellites are dynamic pericentrosomal assemblies that regulate centrosomal protein homeostasis and ciliogenesis. While extensively characterized in vertebrates, their existence and assembly principles in other systems are less well understood. Here, we identify SAS-1 and its interaction partner SSNA-1 as core components of centriolar satellite-like structures in *C. elegans*. We show that SAS-1 satellites occupy a previously unrecognized centrosomal layer positioned between the pericentriolar material and a specialized endoplasmic reticulum. SAS-1 satellites exhibit many features of biomolecular condensates, such as dose-dependent formation, rapid dynamics, and sensitivity to disruption of weak hydrophobic interactions, while their organization depends on the microtubule cytoskeleton. Mechanistically we identify the unstructured and so far, uncharacterized N-terminal region of SAS-1 being functionally important for embryonic viability, satellite formation and binding to microtubules. Our findings demonstrate that *C. elegans* possesses bona fide centriolar satellites that share many features with vertebrate satellites, highlighting their evolutionary conservation and importance across species.

**Graphical Abstract:** 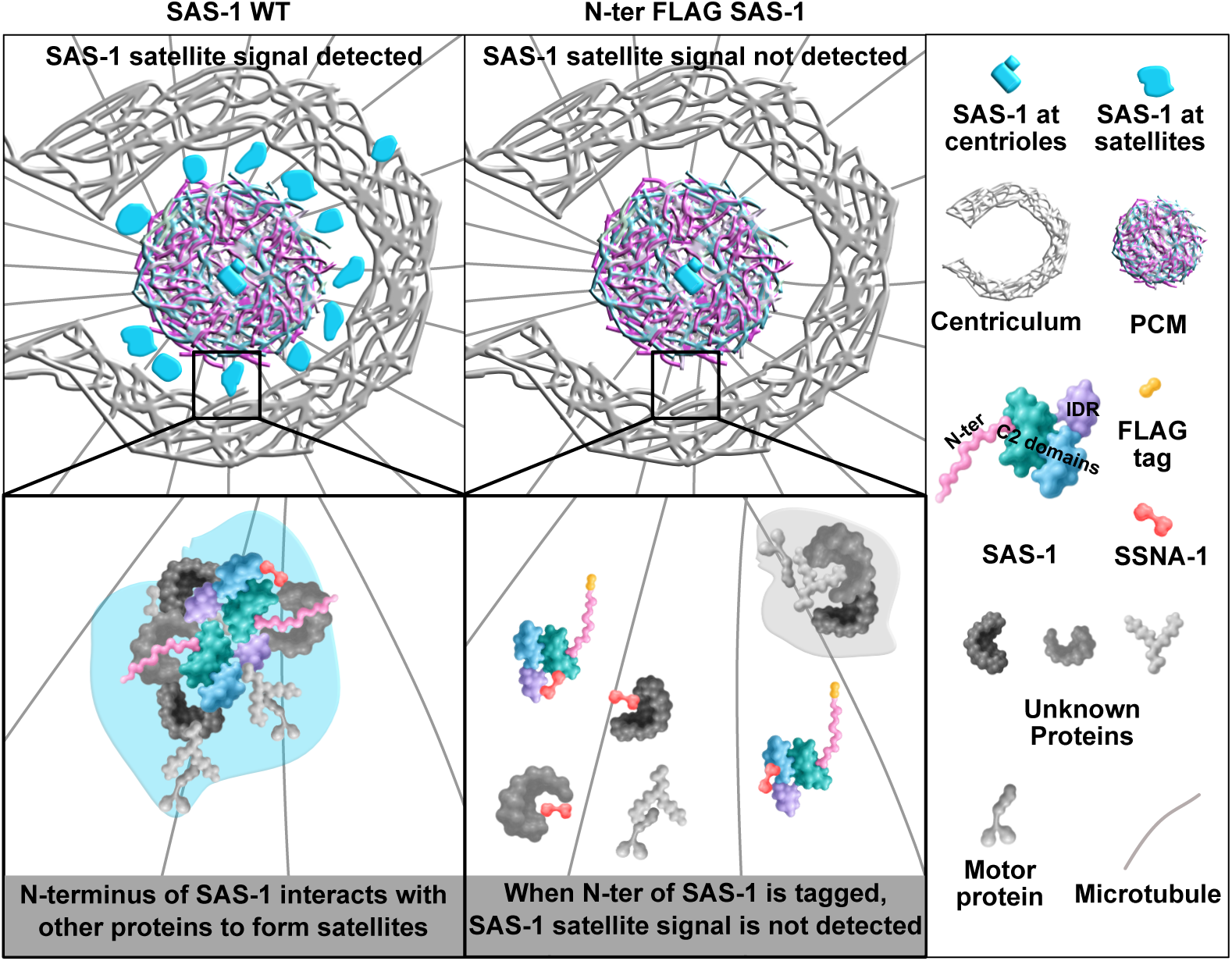

## INTRODUCTION

Centriolar satellites are dynamic membrane-less granular structures, found in the vicinity of centrosomes and cilia. Proteomic studies have identified more than 600 proteins at satellites, many of them overlapping with centrosomal and ciliary proteomes. While the origin and formation of centriolar satellites is not very well understood, their functional relevance in ciliogenesis and neurogenesis is very well documented (reviewed in Begar et al., 2025; Odabasi et al., 2019; Renaud and Bidere, 2021). Centriolar satellites have been implemented in trafficking of centrosome and cilia proteins, proteostasis and autophagy. Most recently, RNA binding proteins and mRNAs of centrosomal proteins (as dSAS4, PCNT and CEP290) were found in close proximity to satellites, suggesting a role in localized protein translation (Pachinger et al., 2025). For many years centriolar satellites were considered to be pericentrosomal structures specific to vertebrate cells, with Pericentriolar material 1 (PCM1) determined as is the main scaffold protein of centriolar satellites in vertebrates. (Dammermann and Merdes, 2002; Kubo et al., 1999; Kubo and Tsukita, 2003). PCM1 is essential for satellite formation and maintenance, as the absence of PCM1 leads to a loss of most centriolar satellites. On the organismic level, PCM1 mutations or depletion in mouse and zebrafish cause ciliopathies and defects in neurogenesis (Hall et al., 2023; Monroe et al., 2020; Odabasi et al., 2019; Stowe et al., 2012). Recently Combover (CMB) was established as the ortholog of PCM1 in *Drosophila melanogaster,* highlighting that centriolar satellites are not exclusively found in vertebrates. Consistent with vertebrate models, downregulation of *cmb* in flies leads to coordination and fertility issues, phenotypes associated with cilia dysfunction (Pachinger et al., 2025). Besides PCM1 being the main scaffold protein and their ciliary function, it is hard to define shared characteristics of satellites across these model systems. Vertebrate satellites are cell cycle-depend, cluster around centrosomes at interphase, and dissolve during mitosis under the influence of dual-specificity tyrosine phosphorylation-regulated kinase 3 (DYRK3) (Rai et al., 2018). Their association with microtubules is dynein- and kinesin-dependent (Dammermann and Merdes, 2002; Kubo et al., 1999; Vicente et al., 2025). Instead, centriolar satellites in flies do not associate with centrosomes and are static, showing no directional mobility along microtubules (Pachinger et al., 2025).

An ortholog or functional homolog of PCM1/CMB has so far not been identified in *Caenorhabditis elegans* (Hodges et al., 2010). Nevertheless, satellite-like structures formed by Sjogren’s Syndrome Nuclear Antigen 1 (SSNA-1) and non-centriolar foci formed by its interaction partner Spindle Assembly −1 (SAS-1) were reported for *C. elegans* (Jha et al., 2025; Pfister et al., 2025). SAS-1 and SSNA-1, and their human homologs C2CD3 and SSNA1, all localize to the distal part of the inner lumen of centrioles and facilitate their structural integrity after formation (Agostini et al., 2025; Bertiaux et al., 2025; Jha et al., 2025; Pfister et al., 2025; Pierron et al., 2023; von Tobel et al., 2014). In addition, C2CD3 is itself a centriolar satellite client protein, supporting the idea that these SAS-1 pericentrosomal foci correspond to centriolar satellites (Ye et al., 2014). However, a detailed analysis of these satellite-like structures was so far not performed. In absence of a reliable ortholog of a scaffold protein and detailed analysis of their properties, it is debatable whether these structures represent *bona fide* centriolar satellites in worms. To close this gap, we set out to explore the dynamic nature of satellite-like structures formed by SAS-1 and SSNA-1 and to establish whether they represent true centriolar satellites in *C. elegans*. We show that SAS-1 and SSNA-1 form centriolar satellites that define a new pericentrosomal layer. Furthermore, we demonstrate that SAS-1 satellites are dynamic, cell cycle and microtubule-dependent and that they exhibit characteristics of biomolecular condensates formed through weak protein-protein interactions. The N-terminus of SAS-1 facilitates centriolar satellite formation and microtubule binding. SAS-1 satellites form in the absence of PCM1 while sharing many features with vertebrate satellites, indicating conserved roles across species.

## RESULTS

### SAS-1 forms dynamic satellite-like pericentrosomal foci

To analyze the localization and dynamics of SAS-1, we generated two endogenously tagged strains with the fluorophore fused to the C-terminus (SAS-1::mkate2 and SAS-1::GFP, Fig. S1A). Live-cell imaging and indirect immunofluorescence staining using both of these strains revealed that SAS-1, as reported previously, localizes to centrioles and at the cilia in the head and tail of the worm (Figs. 1A, S1B, S1C, S1D). In accordance with recent publications, SAS-1 was found at the transition zone, which corresponds to its localization slightly above the GFP::PCMD-1 signal, decorating the ciliary base (Fig. S1D middle and bottom, arrows) (Erpf et al., 2019; Garbrecht et al., 2021; Jha et al., 2025).

**Figure 1.**
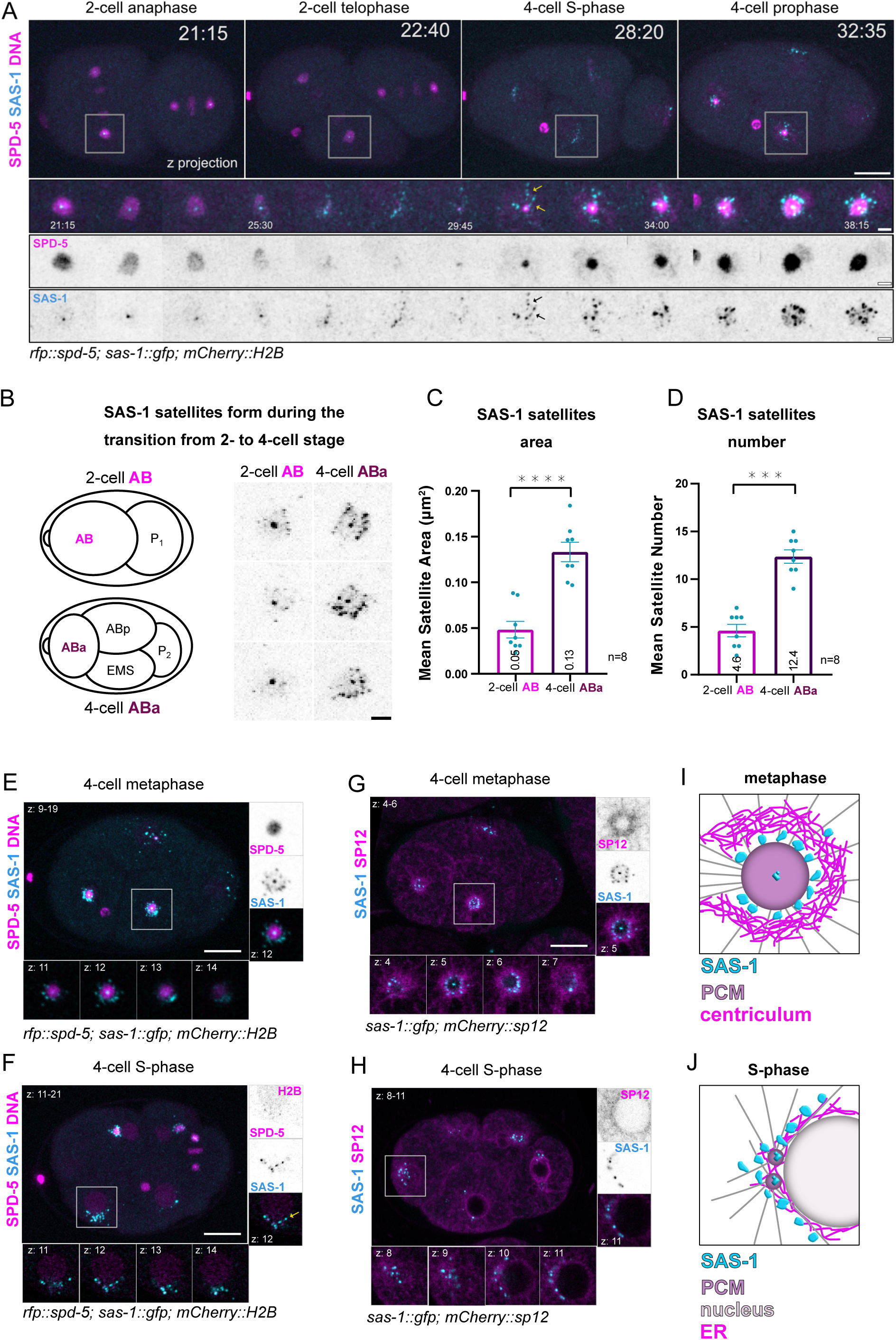
SAS-1 localizes to satellite-like foci. A) Stills of a 2-cell embryo transitioning to a 4-cell stage. SAS-1::GFP first is found at the centriole, gradually forming satellite-like foci, putatively aligning along microtubules (arrows). B) Schematics and enlarged images of SAS-1::GFP at centrosomes in AB (2-cell stage) and ABa (4-cell stage) blastomeres. C) and D) Mean area (C) and number (D) of satellite-like foci around centrosomes in AB (2-cell stage) and ABa (4-cell stage) blastomeres. Significance was assessed using two-tailed paired t-test, error bars represent SEM, n = number of centrosomes analyzed. E) and F) SAS-1 satellite-like foci surround the PCM in metaphase (E) and the nuclear envelope in S-phase (F). G) and H) SAS-1 satellite-like foci line up at the inner rim of the centriculum at metaphase (G) and disperse, surrounding the nuclear envelope in S-phase (H). I) and J) Schematics of SAS-1 satellite-like foci in M- and S-phase. Scale bars 10μm. See also Video 1

Intriguingly, we also noticed that in addition to the centrioles, SAS-1 forms dynamic pericentriolar foci reminiscent of centriolar satellites in early embryos (Fig. 1A). Similar centriolar satellite-like structure has been so far reported for its interaction partner SSNA-1 (Pfister et al., 2025).

To better understand the origin and nature of these foci, we performed live-cell imaging with the PCM marker, RFP::SPD-5, and a histone marker to track cell cycle progression. By live-cell imaging SAS-1 satellite-like foci were not detected in 1-cell embryos, and started appearing at the transition from a 2-cell stage to a 4-cell stage embryo (Fig. 1A, S1C, S1G, Video 1). Satellite-like foci initially become detectable at the prophase when centrosomes start to mature, lining up in the vicinity of the centrioles, seemingly decorating microtubules emanating from the maturing PCM (Fig. 1A, arrows). When the PCM reached its full extent, SAS-1 satellite-like foci gradually arranged around the centrosome, forming a circle around the PCM, when seen in a cross-section. At the transition from a 2-cell to 4-cell embryo, not only did the number of SAS-1 satellite-like foci increase significantly, but also their mean area increased from 50 nm^2^ to 130 nm^2^ (Figs. 1B, 1C, 1D, S1F).

Blastomeres of the *C. elegans* embryos cycle between Mitotic (M)- and Synthesis (S)-phase, without GAP phases. At M-phase, especially metaphase, SAS-1 foci surround the PCM, while in S-phase, after PCM disassembly, satellite-like foci disperse (Figs. 1E, 1F). Interestingly, at S-phase, when the two centrosomes separate and are pushed to opposite sides of the nucleus with the help of microtubules, we also observed a ‘trailing’ of the satellite-like foci along microtubules (Fig. 1F, arrow). SAS-1 satellite-like foci, that did not colocalize with centrioles, were visible in later developmental stages, especially in the vicinity of forming amphid cilia in comma stage embryos (Fig. S1D, top panel, arrow).

The centrosome of *C. elegans* is surrounded by a specialized endoplasmic reticulum (ER), the centriculum (Maheshwari et al., 2023; Maheshwari et al., 2026). The centriculum is a dynamic, cell cycle-dependent structure, thought to act as a siphon that orients microtubules nucleated from the PCM. Since SAS-1 satellite-like structures also surround the PCM, we set out to understand whether they are part of the centriculum. Live-cell imaging of embryos expressing SAS-1::GFP and the ER marker mCherry::SP12 revealed that SAS-1 foci localize at the inner rim of the centriculum in metaphase (Figs. 1G, 1I). Recently, Maheshwari et al., (2023; 2026) reported a gap between the PCM and centriculum, corresponding to the ring of microtubules surrounding the centrosome at mitosis. We speculate that the SAS-1 satellite-like foci are located within this gap. In S-phase, after PCM disassembly, the centriculum disperses which correlates with a more scattered appearance of SAS-1 foci (Fig. 1H, 1J).

Taken together, SAS-1 localizes to dynamic satellite-like structures that first become detectable by live-cell imaging in late 2-cell stage embryos, coinciding with the time of zygotic genome activation. SAS-1 satellite-like structures are sandwiched between the PCM and centriculum at metaphase, defining a so far undescribed pericentrosomal layer. At S-phase, SAS-1 satellite-like structures re-localize to the vicinity of the nuclear envelope (Figs. 1I, 1J).

### N-terminal tagging of SAS-1 interferes with SAS-1 satellite-like foci formation and viability

Although both endogenously tagged strains are fully viable and show no embryonic lethality, as reported for *sas-1* mutant worms (Figs. S1E, 2H), these pericentrosomal foci could still be caused by the presence of a bulky fluorophore (Jha et al., 2025; von Tobel et al., 2014). To verify that this was not the case, we performed immunofluorescent staining on embryos expressing *in situ* N- and C-terminally tagged 3xFLAG::SAS-1 and SAS-1::3xFLAG (Figs. 2A, 2B). Similar to the C-terminally tagged GFP- and mkate2-tagged proteins, SAS-1::3xFLAG satellite-like structures were present most prominently starting at the 4-cell stage blastomere. However, we could not identify centriolar satellites in the N-terminally 3xFLAG::SAS-1 tagged strain (Figs. 2A, 2B). The centriolar signal in 3xFLAG::SAS-1 and SAS-1::3xFLAG was detected. Interestingly, we also noted that in comparison to the C-terminally tagged strain, the N-terminally tagged one 3xFLAG::SAS-1 was less viable, showing only 55% embryonic viability (Fig. 2C). Given that both proteins are highly expressed in whole worm lysates, embryonic lethality, is likely not caused by the reduced expression levels of the N-terminally tagged protein, but rather a functional impairment caused by the N-terminal tag (Fig. 2D). In sum, N-terminal tagging affects SAS-1 localization to satellite-like foci and viability of the embryo.

**Figure 2.**
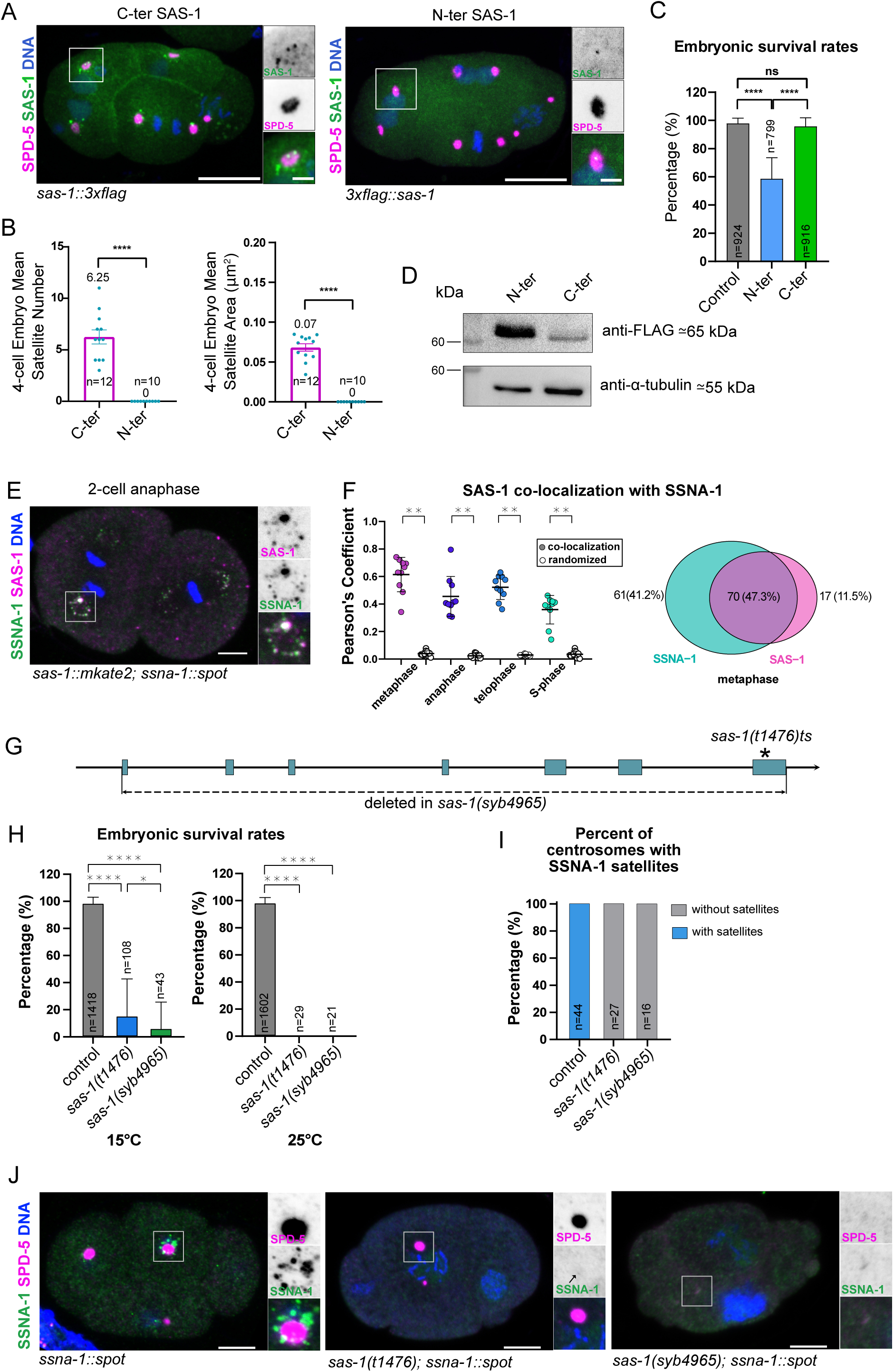
SAS-1 co-localizes with SSNA-1 at satellite-like foci. A) Immunostaining of SAS-1::3xFLAG and 3xFLAG::SAS-1 expressing 4-cell embryo. B) Mean number and area of satellite-like foci around centrosomes in ABa (4-cell stage) blastomeres. Significance was assessed using two-tailed paired t-test, error bars represent SEM, n = number of centrosomes analyzed. C) Embryonic survival rates of control (N2) and SAS-1::3xFLAG and 3xFLAG::SAS-1-tagged strains. Significance was assessed using unpaired Welch’s t-test, error bars are SD n = number of embryos analyzed. D) Western blot of whole worm lysates expressing SAS-1::3xFLAG and 3xFLAG::SAS-1. E) Representative image of a 2-cell embryo stained for SAS-1 and SSNA-1. F) Pearson’s coefficient of colocalization of SAS-1 with SSNA-1 at centriolar satellites at different cell cycle stages and corresponding Venn diagram of overlap. Wilcoxon signed-rank test with Holm correction, error bars represent SD, n = number of foci analyzed. G) Schematic representation of *sas-1* genomic locus and *sas-1(syb4965)* deletion and the point mutation in *sas-1(t1476)ts,* resulting in P419S. H) Embryonic survival rates of control, *sas-1(t1476)ts* and *sas-1(syb4965)* null mutant embryos. Significance was assessed using unpaired Welch’s t-test, error bars are SD, n = number of embryos analyzed. I) SSNA-1::SPOT localization in control, *sas-1(t1476)ts* and *sas-1(syb4965)* null mutant embryos. Arrow points to a weak SSNA-1::SPOT signal at the centrosome. Note that SPD-5 foci are hardly detectable in the *sas-1(syb4965)* null mutants. J) Percent of SPD-5 positive centrosomes showing SSNA-1 satellite-like foci. n = number of centrosomes analyzed Scale bars 10μm.

### SAS-1 satellite-like foci colocalize with SSNA-1

Similar to SAS-1, its interactions partner SSNA-1 also forms satellite-like foci (Pfister et al., 2025). Therefore, we set out to test whether SSNA-1 and SAS-1 localize to the same foci. We crossed SSNA-1::SPOT with SAS-1::mkate2 and quantified their colocalization in immunostaining of embryos at different cell cycle stages using Pearson-correlation analysis (Fig. 2E, F). The majority of the SSNA-1 foci colocalize with SAS-1 satellite-like foci, at all cell cycle stages, indicating that the SAS-1 satellite-like foci harbor at least two proteins. To test whether SAS-1 is required for the SSNA-1 localization to these foci, we used an existing hypermorphic temperature sensitive point mutation *sas-1(t1476)ts* and additionally generated a null allele of *sas-1(syb4965)* by deleting the entire locus of *sas-1* (Fig. 2G). Both *sas-1(t1476)ts and sas-1(syb4965)* are 100% parental embryonic lethal at 25°C; however at 15°C, *sas-1(syb4965)* is significantly less viable than *sas-1(t1476)ts* (Fig. 2D), demonstrating that the null allele is more severe than the point mutation. While we observed a centriolar SSNA-1::SPOT signal at the center of the PCM and satellite-like foci surrounding the PCM in wild type embryos, in *sas-1(t1476)ts* embryos the satellite-like foci were not detected, although a very weak centriolar signal of SSNA-1 was still occasionally present (Figs. 2I arrow, 2J). In *sas-1(syb4965)* embryos we could neither detect a centriolar signal of SSNA-1, nor any satellite-like foci (Figs. 2I, 2J). Interestingly, the *sas-1(syb4965)* null mutant embryos exhibit an extremely weak, almost undetectable SPD-5 signal. This is most likely is due to the structural degeneration of sperm derived centrioles (Jha et al., 2025; von Tobel et al., 2014), and their inability to recruit the PCM and form a microtubule-organizing center (MTOC).

In summary, both SAS-1 and SSNA-1 localize to pericentrosomal satellite-like structures with partial overlap (Pfister et al., 2025). SAS-1 is either required for the formation of these pericentrosomal foci or alternatively, is facilitating SSNA-1 localization to these pericentrosomal foci.

### SAS-1 satellite-like foci follow PCM scaffold dynamics during cell cycle

Since this is the first evidence of centriolar satellites-like structures in *C. elegans* we set out to investigate the nature of these foci and to determine whether they represent *bona fide* centriolar satellites. In cultured cells centriolar satellites are cell cycle dependent – localizing in the vicinity of the centrosome in interphase and dissolving at mitosis (Dammermann and Merdes, 2002; Kubo and Tsukita, 2003). As mentioned above *C. elegans* blastomeres cycle between M- and S-phase, and contrary to mammalian cells, satellite-like structures are visible throughout these cycles. To understand how the distribution of the SAS-1 satellite-like structures dynamically change we set out to quantify the distribution of foci intensity using immunofluorescent staining of blastomeres at different cell cycle stages (Fig. 3A). Given their localization at the pericentrosomal space (Fig. 1A), we were particularly interested to quantitatively assess their position in respect to the PCM scaffold. To measure the distance of SAS-1 satellite-like structures from the center of the centrosome, we used the centriolar SAS-1 signal as the center point, drew 25 concentric circles and plotted total pixel intensities within the rings for SPD-5 and SAS-1 (note that in all graphs the centriolar SAS-1 signal was excluded) (Fig. 3B). We found that together with the PCM scaffold, SAS-1 satellite-like structures follow a canonical pattern. At metaphase SAS-1 intensity peaks at the rim of the PCM, which was defined as the half-max of normalized intensity of SPD-5 (Fig. 3C, left panel). At anaphase the PCM scaffold starts to disassemble by expanding and finally dissolving in S-phase, leaving behind the core PCM surrounding the centrioles. At anaphase, SAS-1 foci are found farther away from the centrioles with a peak located still adjacent to the PCM rim (Fig. 3C, mid panel). However, when the PCM scaffold disintegrates at S-phase, SAS-1 satellites stay more dispersed rather than relocating close to the remaining PCM core (Fig. 3C, right panel).

**Figure 3.**
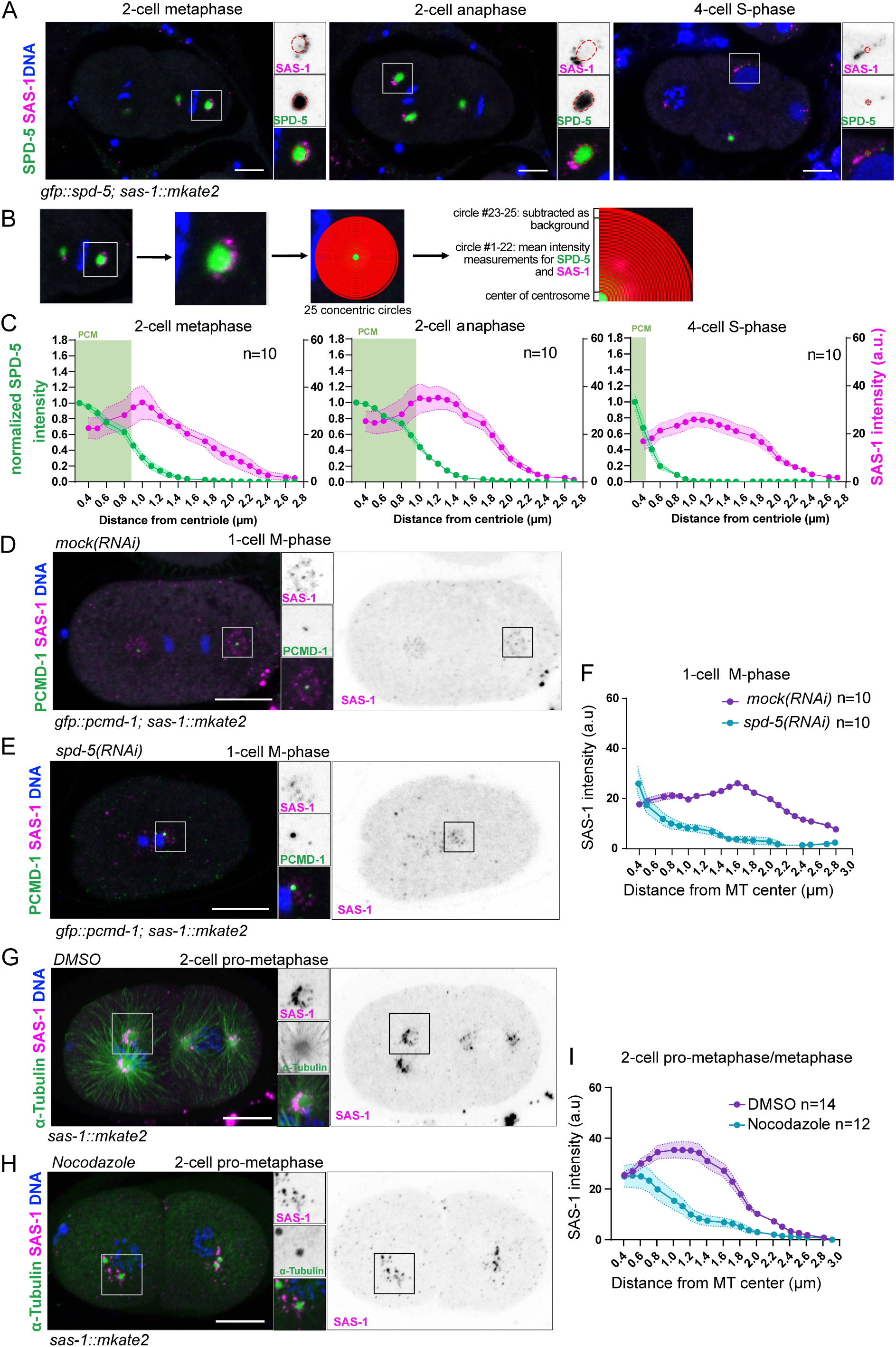
SAS-1 satellite-like foci follow the PCM dynamics during M-phase. A) Representative images of embryos from 2- to 4-cell stained for SAS-1 and SPD-5. B) Images showing the concentric ring analysis to quantify SAS-1 signal intensity around the centrioles. C) Graphs showing SAS-1 signal distribution in comparison to the PCM/centrosome at different cell cycle stages. D) and E) Representative images of with *mock(RNAi)* (D) and *spd-5(RNAi)-*treated (E) embryos stained for SAS-1 and PCMD-1. F) Quantification of SAS-1 distribution in embryos expressing GFP::PCMD-1 and SAS-1::m2ate2 treated with *mock(RNAi)* and *spd-5(RNAi).* G) and H) Representative images of embryos stained with α-Tubulin and SAS-1, treated with DMSO (G) or nocodazole (H). I) Graph showing SAS-1 signal distribution with DMSO or nocodazole treatment. Scale bars 10μm, error bars are SEM, n=number of centrosomes analyzed

Thus, SAS-1 foci are excluded from the PCM and follow the PCM shape during its disassembly but remain distant from the PCM core when the PCM scaffold disintegrates. Therefore, SAS-1 satellite-like structures appear to follow PCM dynamics in M-phase.

To further test this hypothesis, we asked how the spatial distribution of SAS-1 satellite-like structures change in response to manipulations in the PCM size. To do so we used a well-characterized mutation in SZY-20, which is a negative regulator of the centrosome size (Song et al., 2008). When we stained *szy-20(bs52)* mutant and control embryos for SAS-1, satellite-like structures still appeared to be mostly at the border of the enlarged PCM scaffold, with some of them appearing partially embedded in the outer rim (Figs. S2A, S2C). This was confirmed when we plotted the intensity values using the concentric ring analysis and found that in contrast to the control, SAS-1 intensities partially overlapped with the increased PCM diameter in metaphase and anaphase (Figs. S2B, S2D). The PCM is an amorphous structure with changing viscoelastic properties during the cell cycle (Mittasch et al., 2020). The pore diameter of the PCM matrix at metaphase is reported to be less than 15 nm (Tollervey et al., 2025). The measured SAS-1 foci size highly exceeds the pore size, meaning that they would be excluded from the matrix by size exclusion. Perhaps the PCM in *szy-20(bs52)* embryos is not only larger, but also less dense which would enable the SAS-1 foci to integrate into the PCM mesh. To test this hypothesis, we calculated the PCM density in P1 blastomeres using live-cell imaging (Figs. S2E, S2F). Although SAS-1 foci are hardly visible in P1 blastomeres, we specifically chose these cells since the spindle is oriented parallel to the longitudinal axes of the embryo and both spindle poles are located in the same focal plane. At both meta- and anaphase, the PCM area was significantly larger in *szy-20(bs52)* embryos compared to controls (Figs. S2F, S2G, S2H). We calculated overall PCM density by assessing total PCM mass per centrosome volume (Fig. S2E). PCM density is significantly higher at metaphase, and comparable to controls in anaphase in *szy-20(bs52)* embryos (Fig. S2F). Since the PCM density did not decrease this cannot explain the partial overlap of PCM with the SAS-1 intensity. We cannot exclude that rather than the entire PCM scaffold, potentially only the outermost layer might be different in density allowing for the SAS-1 foci to be embedded in it or alternatively, the gap between PCM and centriculum might be narrower or more densely populated with microtubules (Song et al., 2008), pushing the SAS-1 foci into the PCM.

To assess how SAS-1 satellite-like structures respond to the absence of the PCM, we minimized the PCM scaffold by treating embryos with RNAi against SPD-5. Under this condition, the PCM scaffold does not form and only the PCM core remains at the centrosome (Hamill et al., 2002). As a result, the mitotic spindle collapses, cytokinesis fails and cell cycle stages cannot be reliably defined. Therefore, we used the number of centrosomes marked by the PCM core protein PCMD-1 (Erpf et al., 2019; Stenzel et al., 2021), as a proxy to determine the number of the cell cycles and analyzed the distribution of SAS-1 satellites by pooling all M-phase 1-cell embryos, judged by the condensed state of the DNA (Figs. 3D, 3E). While in *mock(RNAi)-*treated embryos SAS-1 satellites were well organized, arranged in a circular manner around the PCMD-1 signal, in *spd-5 RNAi*-treated embryos, they seem to have lost their circular pericentrosomal organization, adopting a dispersed and scattered arrangement, further away from the centrioles, which is reflected by a flattened SAS-1 curve (Figs. 3E, 3F).

Taken together, SAS-1 foci seem to form a pericentrosomal layer filling a narrow gap between the PCM and the centriculum, and following the PCM dynamics during M-phase. In S-phase, after the PCM scaffold disassembles, they lose their pericentrosomal arrangement and appear disperse, without forming a clear peak at the edge of the PCM core.

### Enrichment of SAS-1 satellite-like foci around the PCM is microtubule-dependent

Studies in the 1990s showed that in vertebrate cells centriolar satellites move along microtubules and that their spatial cellular localization is highly dependent on motor proteins (Dammermann and Merdes, 2002; Kubo and Tsukita, 2003; Vicente et al., 2025). When SPD-5 is downregulated by RNAi, less or no microtubules are nucleated from the centrosome (Hamill et al., 2002), and the observed scattered distribution of the SAS-1 satellites could be due to the lack of microtubule nucleation in absence of a PCM scaffold. Could SAS-1 satellites be transported along microtubules? During live-cell imaging we observed that some of the SAS-1 satellite-like structures formed characteristic trails (Figs. 1A, 1F), suggesting an alignment along microtubules emanating from the centrosomes.

To test how SAS-1 foci behave in absence of microtubules, we depolymerized microtubules without affecting the MTOC. To do so, we permeabilized the egg shell by using *perm-1(RNAi)* and acutely depolymerized microtubules by exposure to nocodazole for 3 min before fixation. Successful microtubule depolymerization was monitored by staining with an antibody against α-Tubulin (Figs. 3G, 3H). After acute microtubule loss, majority of SAS-1 foci were observed to lose their typical circular arrangement and enrichment around the PCM at a distance of 0.8 μm – 2 μm (Fig. 3I). The SAS-1 intensity peak appears to be lower, indicating that overall less SAS-1 foci accumulated next to the centrosome, with some remaining foci clustering in closer vicinity to the centrosome, at 0.4 μm-0.8 μm away from the center. Additionally, we found a population of foci dispersed throughout the blastomeres, farther away from the radius usually measured by the concentric rings (Figs. 3H, S3A, S3B).

Similar to nocodazole, cold treatment can be used to depolymerize microtubules. We incubated embryos for 20 min on a cooled block before fixing them for immunofluorescence staining. In response to cold-treatment, microtubules depolymerize, leading to the collapse of the mitotic spindle (Figs. S3C, S3D, S3E). As in nocodazole-treated embryos SAS-1 foci appeared closer to the PCM. To quantify the precise localization of SAS-1 satellites in respect to the PCM scaffold, we used SPD-5 as a marker and performed concentric ring analysis on cold-treated embryos (Figs. S3E, S3F). When embryos were cold-treated, SAS-1 foci seemed to coalesce in close vicinity to the PCM scaffold. Note that some of the foci occasionally appeared fused into larger foci (Fig. S3D, arrow).

Overall, SAS-1 foci distribution in nocodazole- and cold-treated embryos differs from the *spd-5(RNAi)-*treated ones. However, this can be explained by the difference in the treatment. *spd-5(RNAi)* is a systemic treatment, where the PCM scaffold is never formed and no microtubules are nucleated from the beginning of embryogenesis. If SAS-1 satellite-like structures would be microtubule-associated and transported along microtubules, they would never be able to associate with microtubules. During nocodazole and cold treatment, SAS-1 satellite-like structures that are already associated with microtubules are assayed for their dislocation upon depolymerization of the microtubules. One also has to take in consideration that the absence of microtubules strongly affects the centriculum (Maheshwari et al., 2023). In nocodazole-treated embryos the centriculum is absent, while the centrosome is still associated with endoplasmic reticulum surrounding the nuclear envelope. If SAS-1 foci would be confined by the centriculum, we would expect their dispersal upon microtubule depolymerization. Their close proximity to the PCM indicates a dependence on microtubules, rather than centriculum. Overall, we conclude that SAS-1 satellite-like structures change their localization in response to microtubule depolymerization and are therefore microtubule-dependent.

### SAS-1 satellite-like foci are dynamic structures

We next decided to examine dynamic properties of SAS-1 at the centrosome. To assess whether there is a difference in SAS-1 dynamics between the satellite-like and centriolar pools, we performed fluorescence recovery after photobleaching (FRAP) on 2-cell embryos in late telophase, when satellite-like structures start to be detectable by live-cell imaging (Fig. 4A, Video 2). At one spindle pole, the entire volume of one centrosome and its surroundings was bleached, while the second pole was used as an internal control. Recovery of SAS-1 signal at the centrioles and satellites was quantified independently.

**Figure 4.**
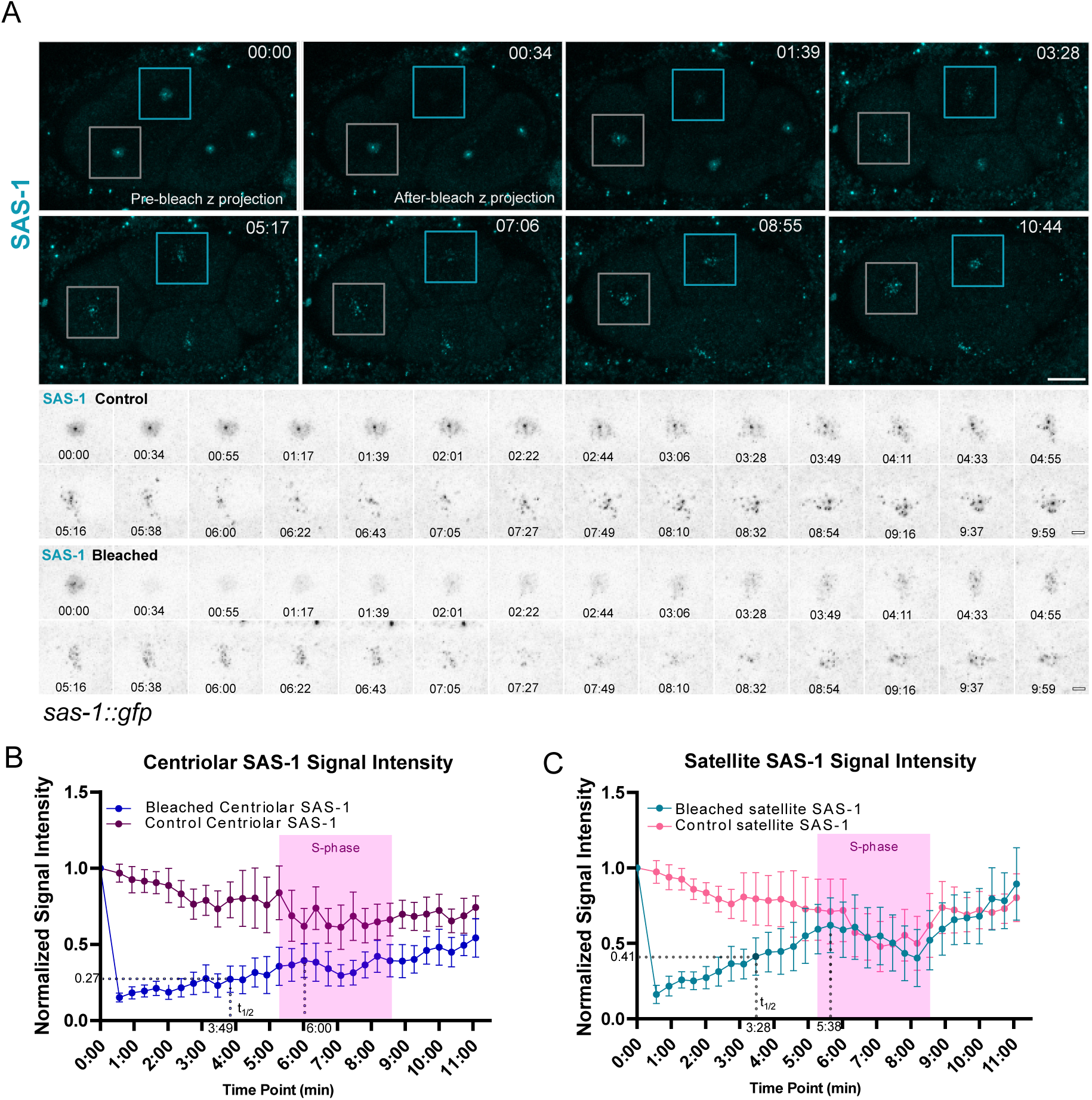
The SAS-1 pool at satellite-like foci is more dynamic than at centriolar pool. A) Stills from a movie showing FRAP of one spindle pole in AB cell, while the second spindle pole serves as an internal control. B) Normalized SAS-1 signal intensities at the centrioles with and without FRAP. C) Normalized SAS-1 signal intensities at satellites with and without FRAP. Scale bars 10μm and 2μm in insets, error bars are SEM, n=number of embryos analyzed See also Video 2

The centriolar SAS-1 pool showed very limited mobility and a slow and incomplete recovery kinetics (with 38% recovery plateau and ½ time of 3 min 49 sec) (Fig. 4B). After the cells passed through S-phase the signal exceeded its 50% of its original intensity. This can be explained by the formation of a new daughter centriole that takes place during S-phase. Overall, this indicates that the SAS-1 pool at centrioles mostly consists of a non-dynamic fraction, which is consistent with its role in centrioles stability.

Instead, the satellite pool recovered relatively fast. First a diffuse cloud became visible, from which small SAS-1 foci start appearing over time. The bleached pool recovered to a maximum of 60% with a ½ time of 3 min 28 sec. Note that due to the dispersal of satellites at S-phase, the values of the control pool dropped as well and were similar to the bleached pool at the timepoint of its maximal recovery (Fig. 4C). After this timepoint both curves follow the same dynamics. These results argue for a dynamic nature of SAS-1 at satellite-like foci.

### SAS-1 satellite-like foci require hydrophobic interactions

Very little is known about the molecular mechanism of centriolar satellites formation. Recently it was suggested that newly formed centriolar satellites are sensitive to 1,6-hexanediol (1,6-HD) (Begar et al., 2026), a reagent that disrupts weak hydrophobic interactions and is frequently used to probe dynamic states of membraneless compartments (Putnam et al., 2019; Updike et al., 2011). This is in line with previous reports that the activity of DYRK3 promotes dissolution of satellites at mitosis ((Rai et al., 2018). Both observations suggest that centriolar satellites could form as biomolecular condensates.

SAS-1 satellite-like foci become visible in the two-cell embryos coinciding with the activation of the zygotic genome, suggesting that their formation may be concentration dependent, a property often exhibited by biomolecular condensates. To test whether SAS-1 satellites are formed by weak hydrophobic interactions, we permeabilized embryos using *perm-1(RNAi)* and exposed them to 1,6-HD (Figs. 5A and 5B, Videos 3 and 4). Four-cell stage embryos were imaged for the duration of 8 minutes after adding 1,6-HD. We quantified separately the PCM scaffold and the SAS-1 pool at the centrioles and satellite-like foci. After 1,6-HD exposure, RFP::SPD-5 signal rapidly diminished (Figs. 5B and 5D). This is interesting, since SPD-5 itself is a protein that undergoes liquid-liquid phase separation (LLPS) *in vitro* (Woodruff et al., 2017; Woodruff et al., 2015), however it is debatable whether the PCM has properties of a biomolecular condensate (Raff, 2019). Similar to SPD-5, the pool of SAS-1 at the satellite-like foci was completely lost after a few minutes of drug treatment. In contrary, SAS-1 at the centrioles diminished only marginally, acting as a positive control during imaging (Figs. 5A and 5C).

**Figure 5.**
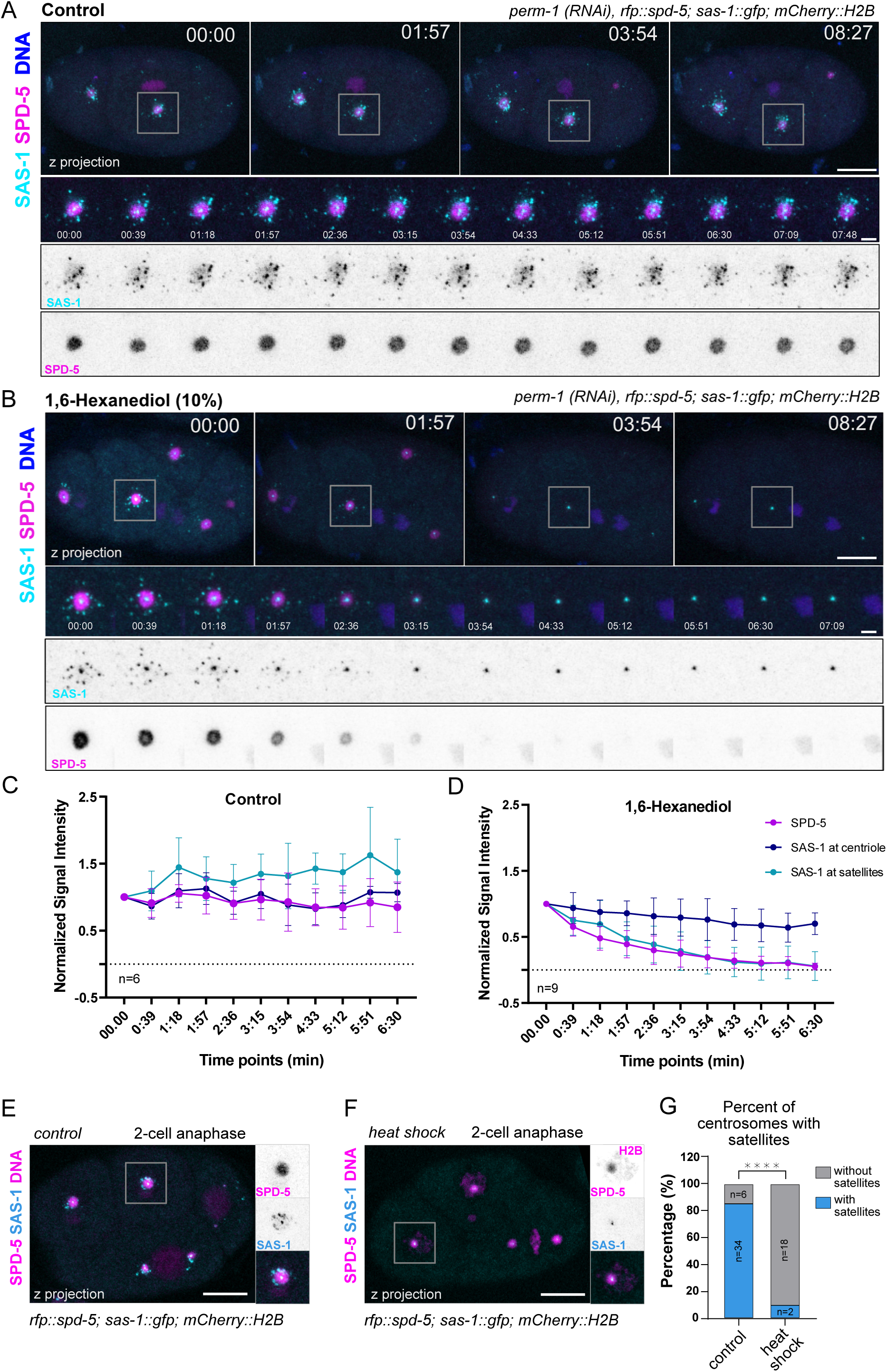
SAS-1 satellite-like foci require hydrophobic interactions for their maintenance. A) Stills of *perm-1(RNAi)-*treated control embryos, expressing SAS-1::GFP. B) Stills of the same embryos treated with 10% 1,6-Hexanediol. Note: PCM scaffold and satellite-like foci are lost, while SAS-1 at the centrioles is not affected. C) and D) Quantification of RFP::SPD-5, SAS-1::GFP at centrioles and satellites in control and 1,6-Hexanediol-terated embryos. n=number of embryos analyzed. E) and F) Imaged of 4-cell embryos expressing RFP::SPD-5 and SAS-1::GFP without (E) and with (F) exposure to one hour heat-shock of 34°C. G) Quantification of centrosomes with and without SAS-1::GFP at satellites. Significance was assessed using two-sided Fisher’s exact test, n=number of centrosomes analyzed. Scale bars 10μm and 2μm in insets; error bars are SEM See also Video 3 (control) and Video 4 (1,6-Hexanediol treatment)

Biomolecular condensates often form under specific environmental conditions and are susceptible to abiotic stress. A good example are P-granules in the *C. elegans* embryos and germline, which reversibly dissolve when the animals are treated with heat shock at 34°C for one hour (Putnam et al., 2019; Watkins and Schisa, 2021). We reasoned that SAS-1 foci could be affected by a similar heat stress. Indeed, after incubating embryos for one hour at 34°C, we could not observe any SAS-1 satellites at the centrosomes (Figs. 5E, 5F, 5G), while SAS-1 at the centrioles was still detectable. Intriguingly, the PCM scaffold that dissolved under 1,6-HD treatment, remained largely resistant to the abiotic heat stress.

Taken together we conclude that the two pools of SAS-1 not only have different dynamic properties, but are formed by different kinds of interactions, where the pool at the satellites depends on weak hydrophobic interactions sensitive to 1,6-HD treatment and heat stress.

### SAS-1 satellites formation is indeed concentration-dependent

If SAS-1 satellites formation is indeed concentration-dependent, we should be able to observe earlier satellite formation when SAS-1 is overexpressed. To this end we generated a C-terminally tagged SAS-1::GFP and an N-terminally tagged GFP::SAS-1 transgene integrated as a single copy and driven by the *mai-2* regulatory elements (Fig. 6A). Both strains showed higher expression in comparison to the endogenous CRISPR-tagged SAS-1::GFP (Fig. 6B). We could clearly identify pmai-2:SAS-1::GFP satellites forming earlier, already in one-cell stage embryos with, while the endogenously-tagged SAS-1::GFP foci were hardly visible at the same timepoint (Figs. 6C, 6D, Video 5). Satellites formed by the transgene appear not only earlier, however are higher in numbers and larger (Figs. 6F, 6G). Interestingly, at 4-cell stage, in the overexpression strain satellite-like foci seemed to fuse or cluster (Fig. S4A, S4B, S4C, S4D). Despite its high levels of expression, relatively few foci were formed by the N-terminally tagged pmai-2:GFP::SAS-1 in the presence of the endogenous protein (Figs. 6E, 6F, 6G). This is similar to the absence of centriolar satellites in the endogenously N-terminally-tagged 3xFLAG strain (Figs. 2A and 2C). Together with the increased lethality of the 3xFLAG::SAS-1 strain, this data suggests a functional relevance of the N-terminus for SAS-1 function, which could be due to the lack of satellite formation or SAS-1 localization to the same.

**Figure 6.**
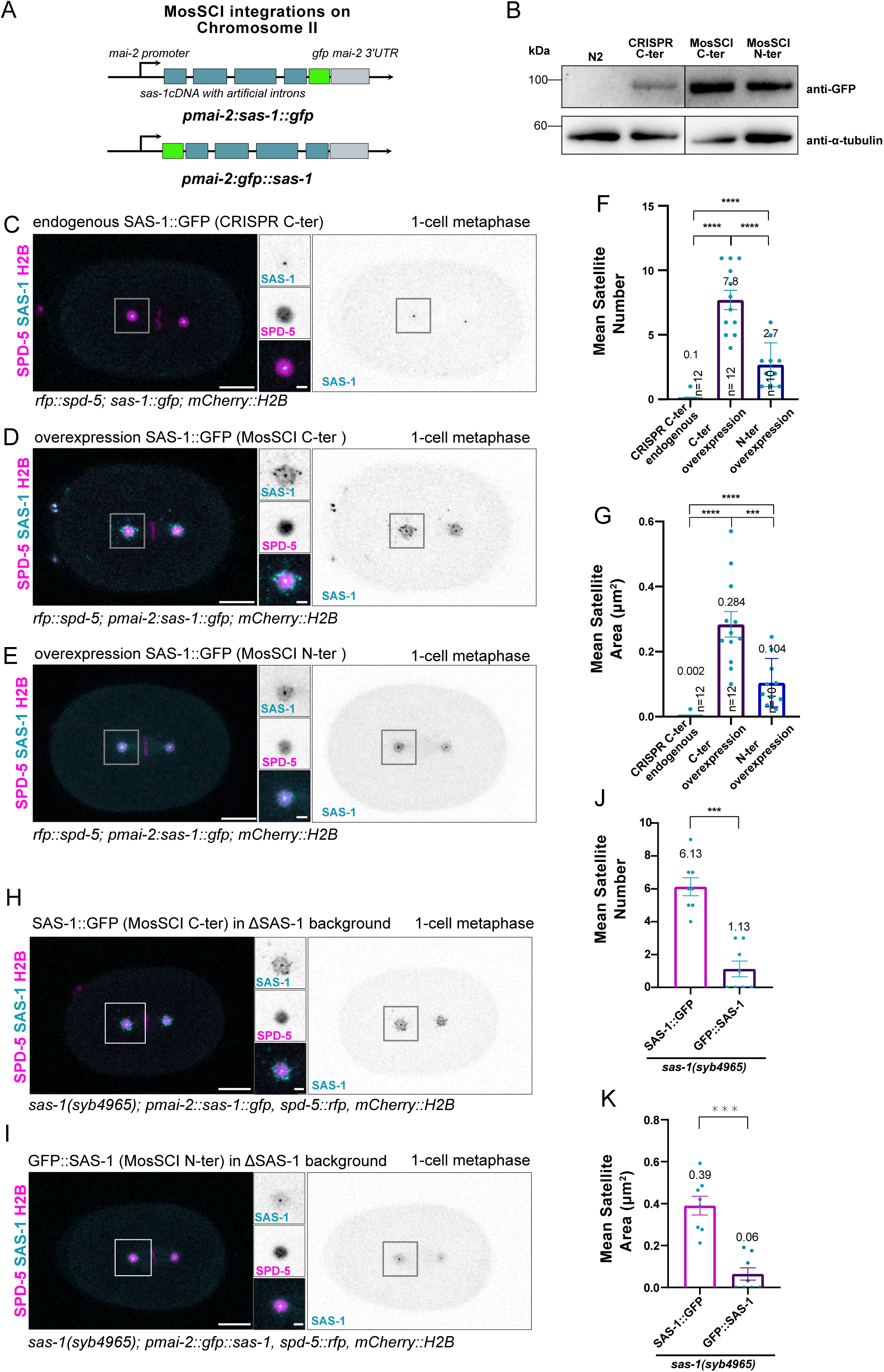
SAS-1 satellite-like structures form in a dose-deepened manner. A) Schematic representation of the C-terminally and N-terminally GFP-tagged MosSCI transgene. B) Immunoblot showing expression levels of endogenously GFP-tagged, C-terminally GFP-tagged SAS-1 and N-terminally GFP-tagged SAS-1 overexpression strains along with N2 strain shown as negative control. The images are from the same immunoblot. C) Stills of a one-cell embryo expressing endogenously tagged SAS-1::GFP at metaphase. D) Stills of pmai-1::SAS-1:GFP in one cell-embryo at metaphase. E) Stills of pmai-1::GFP::SAS-1 in one cell-embryo at metaphase. F) Mean number of satellite-like foci at centrosomes in one-cell embryos. G) Mean area of satellite-like foci at centrosomes in one-cell embryos. H) Stills of a one-cell embryo expressing pmai-1::SAS-1:GFP in the *sas-1(syb4965)* null mutant background at metaphase. I) Stills of a one-cell embryo expressing pmai-1::GFP::SAS-1 in the *sas-1(syb4965)* null mutant background at metaphase. J) Mean number of satellite-like foci at centrosomes in one-cell embryos. K) Mean area of satellite-like foci at centrosomes in one-cell embryos. Scale bars 10μm, significance was assessed using Mann-Whitney-U test for satellite number and area analysis, error bars are SEM, n = number of centrosomes analyzed. See also Video 5

To test the ability of the two different transgenes in SAS-1 satellite formation in absence of the endogenous protein, we crossed both pmai-2:GFP::SAS-1 and pmai-2:SAS-1::GFP into the *sas-1 (syb4965)* null mutant background (Figs 6H, 6I). In contrast to the C-terminally tagged pmai-2:SAS::GFP transgene, almost no foci be detected in one cell embryos in the N-terminally tagged pmai-2:SAS-1::GFP strain, indicating that the endogenous SAS-1 copy facilitated satellites formation (Figs 6J, and 6K).

Thus, SAS-1 satellite-like structure formation is indeed dose dependent, consistent with a biomolecular condensate-like nature of the SAS-1 foci. N-terminal tagging interferes with either SAS-1 satellite formation or SAS-1 localization to the satellites in the *C. elegans* zygote.

### The N-terminus of SAS-1 is required for its localization to the microtubules

Intrigued by these findings we set out to test the function of the N-terminus of SAS-1. Previous studies hypothesized that the N-terminus of SAS-1 is needed for microtubule binding, while the C-terminus interacts with SSNA-1 (Jha et al., 2025), however the involvement of the N-terminus in microtubule binding was formally not tested. When expressed in mammalian cells, SAS-1 is targeted to stable microtubules (von Tobel et al., 2014). To investigate the role of the N-terminus of SAS-1 in MT-targeting we used a similar cell culture system. The N-terminus of SAS-1 contains one of the two C2 domains (Figs. 7A and 7B). We C-terminally EGFP-tagged either the full-length SAS-1 or SAS-1 in which we deleted the first 31 amino acids (SAS-1(Δ2-31)) or the C2-1 domain (SAS-1(Δ32-184)), and transiently expressed these transgenes in HeLa cells (Fig. 7B and 7C). As reported previously full-length SAS-1::EGFP localizes to microtubules. SAS-1-decorated microtubules appear more bundled and stabilized, than the ones in cells not expressing SAS-1. Interestingly, when we expressed SAS-1(Δ2-31)::EGFP, lacking the most N-terminal 31 amino acids, microtubule localization was abrogated (Figs. 6G and 6H). Instead, SAS-1 diffused in the cytoplasm and formed foci of different sizes with well-defined outlines, reminiscent of protein condensates. Microtubule localization was also abrogated in SAS-1 lacking the C2-1 domain, SAS-1(Δ32-184)::EGFP (Figs. 7C and 7D). However, SAS-1(Δ32-184)::EGFP was not detected in the cytoplasm and formed more prominent protein structures, with more irregular borders reminiscent of protein aggregates rather than condensates (Figs. 7C and 7D). These structures localized to the vicinity of the nuclear envelope, excluding microtubules. To test whether in the absence of microtubule binding SAS-1 is prone to form condensates that are based on weak hydrophobic interactions, we treated the cells with 1,6-HD. While the presence of SAS-1(Δ2-31)::EGFP foci was highly reduced in number, structures formed by SAS-1(Δ32-184)::EGFP were unaffected (Figs. 7C and 7E). This indicates that the foci formed by SAS-1(Δ32-184)::EGFP are qualitatively different, resembling stable protein aggregates.

**Figure 7.**
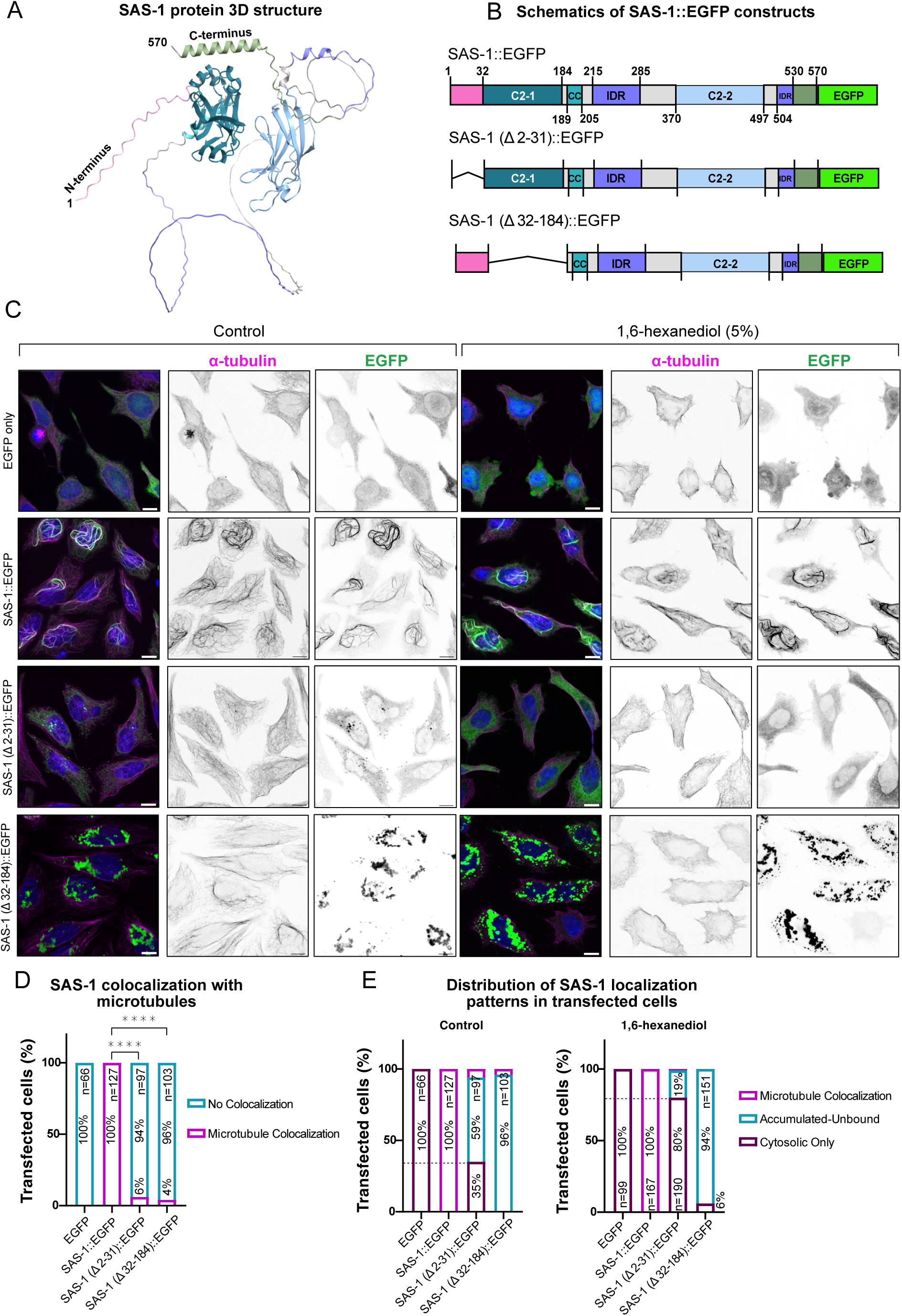
The SAS-1 N-terminal region is required for microtubule localization. (A) Predicted three-dimensional structure of full-length SAS-1, with the N- and C-terminus indicated. (B) Schematic representation of SAS-1::EGFP constructs used in this study. Full-length SAS-1::EGFP contains the N-terminal region, C2-1 domain, coiled-coil region, intrinsically disordered regions, C2-2 domain, and the C-terminal EGFP tag. Deletion constructs lack either residues 2–31 or residues 32–184. (C) Representative fluorescence images of HeLa cells, expressing EGFP alone, full-length SAS-1::EGFP, SAS-1(Δ2–31)::EGFP, or SAS-1(Δ32–184)::EGFP under control conditions or following treatment with 5% 1,6-Hexanediol. Cells were stained for α-tubulin, and the GFP signal was used to assess SAS-1 construct localization. Full-length SAS-1::EGFP colocalized with microtubules, whereas SAS-1(Δ2–31)::EGFP showed reduced microtubule association. SAS-1(Δ32–184)::EGFP formed prominent accumulated/unbound cytoplasmic structures that did not colocalize with microtubules. Scale bar shows 10μm. (D) Quantification of transfected cells classified as showing microtubule colocalization or no colocalization for each construct. Numbers above bars indicate the number of transfected cells analyzed. Localization distributions differed significantly between constructs: Pairwise comparisons were performed using two-tailed Fisher’s exact tests with correction for multiple comparisons; ****P < 0.0001; ns, not significant. (E) Distribution of localization phenotypes among transfected cells expressing SAS-1::EGFP constructs under control conditions or following treatment with 5% 1,6-Hexanediol. Cells were classified as microtubule-bound, accumulated-unbound, or cytosolic only. Full-length SAS-1::EGFP was 100% microtubule-bound in all transfected cells, whereas SAS-1(Δ2–31)::EGFP was predominantly cytosolic with a large subset of cells showing accumulated foci, and SAS-1(Δ32–184)::EGFP predominantly displayed 96% accumulated/unbound localization. Following 1,6-Hexanediol treatment, SAS-1(Δ32–184)::EGFP accumulations persisted, whereas SAS-1(Δ2–31)::EGFP foci accumulation were reduced, with a corresponding shift toward the cytosolic-only phenotype, consistent with dispersal.

Taken together, we conclude that the N-terminal part of SAS-1 is needed to target the protein to the microtubules and tags positioned at the N-terminus might interfere with its function.

## Discussion

Our systematic analysis indicates that besides centrioles SAS-1 localizes to dynamic satellite-like structures. SAS-1 satellite-like foci exhibit many properties of centriolar satellites. They are positioned around the centrosome in a cell cycle and microtubule-dependent manner and are dynamic in nature. SAS-1 satellite-like foci are occupying a defined pericentrosomal space at metaphase and disperse after PCM disassembly at mitotic exit. At metaphase they are sandwiched in a narrow gap between the PCM and the centriculum, a specialized ER, surrounding the centrosome. *C. elegans* PCM proteins occupy different domains, forming layered concentric spheres around the centrioles (Magescas et al., 2019; Mittasch et al., 2020). All PCM proteins analyzed, except of the Aurora kinase-1 (AIR-1) overlap with the PCM scaffold, defined by SPD-5. AIR-1 only partially overlaps with SPD-5, extending beyond its perimeter. Instead, SAS-1 foci border the SPD-5 scaffold, and thereby define a so far undescribed pericentrosomal layer. SAS-1 satellite-like foci could act as a buffering reservoir, regulating protein homeostasis of SAS-1 and perhaps other centrosomal proteins. One of these proteins could be SSNA-1, as both SAS-1 and SSNA-1 localize to these pericentrosomal satellite-like structures with partial overlap (Pfister et al., 2025). We show that SAS-1 is either required for the formation of these pericentrosomal foci or alternatively, facilitates SSNA-1 recruitment. Pfister et al. (2025) also reported that SAS-1 localizes to centriole satellite-like structures in a SSNA-1-dependent manner. Therefore, either SSNA-1 and SAS-1 together form satellite-like foci or similar to centrioles, they are interdependent in their localization to satellites. To discriminate between these possibilities a third satellite marker must be identified in the future.

While centriolar satellites in mammalian systems are microtubule-dependent, in *Drosophila* they are static, showing no directional mobility along microtubules (Dammermann and Merdes, 2002; Kubo et al., 1999; Pachinger et al., 2025; Vicente et al., 2025). Similar to mammalian systems, we found that SAS-1 satellite-like foci seem to align along microtubules, and change position upon microtubule depolymerization. We therefore conclude that they are microtubule-dependent. Although our data demonstrate microtubule dependence, they do not yet establish whether SAS-1 satellites undergo active motor-driven transport or are passively organized by the microtubule network.

Tagging SAS-1 endogenously with a small 3xFLAG-tag at the N-terminus interferes with either SAS-1 localization to satellite-like foci or even formation of these structures. This is in stark contrast to the C-terminally 3xFLAG- tagged SAS-1 where satellite-like foci were detected throughout embryogenesis. Since SSNA-1 was predicted to bind the C-terminus of SAS-1 (Jha et al., 2025), we speculate that the N-terminus is needed for SAS-1 localization to satellites or their formation via interactions with other yet unknown protein. When the N-terminus is carrying the 3xFLAG-tag, these interaction sites are blocked, hindering SAS-1 from localization to satellites. Interestingly, the same N-terminally but not C-terminally 3xFLAG-tagged SAS-1 strain exhibits high embryonic lethality underscoring the importance of the N-terminus. We found that 3xFLAG::SAS-1 protein levels are not reduced but rather elevated and that the protein still localizes to centrioles. Similarly, Woglar et al. reported no ultrastructural abnormalities of centrioles in animals expressing the same 3xFLAG::SAS-1 protein (Woglar et al., 2022). Taken together, these findings indicate that N-terminal tagging does not destabilize the SAS-1 protein and does not affect its centriolar localization, or centriole ultrastructure. Therefore, we favor the hypothesis that the satellite-associated pool of SAS-1 contributes to embryonic viability, independently of its centriolar structural role.

Interestingly, similar to SSNA-1, SAS-1 has the ability to bind microtubules (Agostini et al., 2025; von Tobel et al., 2014). When expressed in HeLa cells, SAS-1 localized to microtubules and seems to bundle them, and we could show that the most N-terminal 31 unstructured amino acids are absolutely necessary for this localization. When this part of the protein is deleted, SAS-1 is unable to bind microtubules, and forms droplets in the cytoplasm. In *C. elegans* embryos microtubules are not necessary for satellites formation but influence their distribution. This is different to the N-terminal tagged SAS-1 strain where no satellites are observed. Therefore, we favor the view that microtubules play a modulatory role, rather than being directly involved in their formation.

Supporting the potential dichotomy of SAS-1 function at the centrioles and satellite-like foci, we found that the SAS-1 pool at centrioles behaves differently from the one at pericentrosomal foci. Firstly, we found that SAS-1 at satellite-like foci is more dynamic than the one at centrioles. Secondly, their formation appears to rely on a distinct set of molecular interactions. While the pool at the satellite-like foci depends on weak hydrophobic interactions, sensitive to 1,6-HD treatment and heat stress, the centriolar, less dynamic pool of SAS-1 is not affected by these treatments. Similar to the centriolar satellite pool, when SAS-1 is expressed in cell culture and delocalized from microtubules by deletion of the unstructured N-terminus, it forms condensates that are sensitive to 1,6-HD treatment. Solubility of these condensates seems to depend on the first C2 domain, in the absence of which SAS-1 forms aggregates, insensitive to 1,6-HD treatment. Consistent with a condensate-like behavior, increasing SAS-1::GFP levels using a strong promotor, reduced the number of satellite-like foci while significantly increasing their area, suggesting that higher SAS-1 concentration promotes coalescence or fusion of foci. Overall, we propose that SAS-1 satellite-like foci are dynamic and sensitive to agents disrupting weak hydrophobic interactions, consistent with a model in which SAS-1 satellites form as biomolecular condensates.

Interestingly, we also observed that that the PCM scaffold protein SPD-5 is rapidly dissolved under 1,6-HD treatment. While this is not a proof that the centrosome has properties of a biomolecular condensate, to our knowledge it is the first evidence that the PCM is sensitive to agents disrupting weak hydrophobic interactions. We do not expect that the PCM scaffold and the SAS-1 satellite-like foci are mutually dependent, although we cannot formally exclude this possibility.

In summary, we propose that SAS-1 together with SSNA-1 forms *bona fide* centriolar satellites in *C. elegans* despite the absence of an identifiable PCM1/CMB homolog, highlighting their evolutionary conservation and importance across species. We establish that the N-terminus of SAS-1 is needed for centriolar satellite formation and embryonic development. SAS-1 satellites exhibit many features of biomolecular condensates, including dose-dependent formation, rapid dynamics, and sensitivity to disruption of weak hydrophobic interactions. Our work opens new avenues for future mechanistic studies using *C. elegans* as a powerful developmental model organism. Future studies will be required to determine whether proteins other than SSNA-1 and SAS-1 are associated with these satellites, and whether a scaffolding protein functionally replacing PCM1/Cmb is present in *C. elegans*.

## MATERIALS AND METHODS

### *C. elegans* strains and culture conditions

*C. elegans* strains were maintained under standard conditions on nematode growth medium (NGM) plates seeded with *Escherichia coli* OP50 strain and kept at 15°C unless otherwise indicated (Brenner, 1974). The Bristol N2 strain was used as wild-type control. All strains used in this study are listed in Supplemental Table 1. For experimental assays, *sas-1(t1476)* homozygous mutant worms were shifted to the restrictive temperature of 25°C at the late L3 or L4 larval stage and incubated for 16-20 hours prior to analysis. For *szy-20(bs52)* mutant, worms were shifted to restrictive temperature of 25°C and incubated for 20 hours before immunofluorescence staining or live-cell imaging. Heat-shock analysis was performed by incubating young adult worms at 34°C for 1 hour. Following heat-shock, worms were dissected, mounted and subjected to live-cell imaging at 25°C.

### Worm strain generation

The *sas-1(syb4965)* null allele was generated by SunyBiotech. This allele carries a 11,062bp deletion within Y111B2A.24, flanked by ttcaaaattttaaaacttctttcagaacta – ttattttttttatttcttattacaactttt, removing the entire coding sequence.

Transgenic worm strains were generated by single-copy integration using the MosSCI system (Frokjaer-Jensen et al., 2012; Frokjaer-Jensen et al., 2008). Transgenes were cloned into the pCFJ350 vector and injected into the EG6699 (chromosome II).

### RNA-mediated interference (RNAi)

RNAi by feeding was performed by placing L4 larvae onto plates seeded with mock or *spd-5(RNAi)* bacteria (clones I-4O08 and I-2G08, respectively), and by incubating them at 25°C for 16-20 hours. For eggshell permeabilization, L4-stage worms were picked the day prior to imaging, transferred to NGM plates seeded with *perm-1(RNAi)* (clone II-5J22) bacteria, and incubated for 20 hours at 20°C (Carvalho et al., 2011).

### Indirect Immunofluorescence

For immunofluorescence (IF) assays*, C. elegans* embryos were isolated by dissecting gravid adults on poly-L-lysine-coated slides. Eggshell was opened by freeze cracking. Samples were fixed in ice-cold methanol for 10 minutes and blocked in 2% BSA for 15 minutes. Slides were incubated with primary antibodies overnight at 4°C, followed by secondary antibodies with Hoechst for 90 minutes at room temperature, DNA condensation patterns were used to stage embryos. For HeLa IF assays, 150.000 HeLa cells were seeded on 6-well plate on a coverslip, in 2mL complete DMEM medium. Cells were transfected for 24 hours, washed with PBS, and fixed with ice-cold methanol for 7 minutes. Cells were blocked for one hour at room temperature then incubated in primary antibody solution for one hour at room temperature. After the washing steps, slides were incubated in secondary antibody solution for one hour. The following antibodies raised in rabbits were used at the indicated concentrations: 1:500 anti-RFP (#600-401-379, Rockland), 1:300 anti-GFP (#A11122, Invitrogen), and 1:1000 anti-SPD-5 (generous gift from B. Bowerman). The following primary antibodies raised in mouse were utilized: 1:500 anti-GFP (#11814460001, Sigma), 1:500/1:300 anti-α-tubulin (DM1α #T6199, Sigma), 1:500 anti-FLAG (#F1804, Sigma) and 1:500 anti-SPOT (#28a5, Chromotek). For secondary antibodies, goat anti-mouse coupled to Alexa488 (#A11001, Invitrogen), goat anti-rabbit coupled to Alexa568 (#A11011, Invitrogen), anti-rabbit Alexa488 (#A11034, Invitrogen) and anti-mouse Alexa594 (#A21203, Invitrogen), all used at 1:500. Slides were counterstained with 1 mg/ml Hoechst33258 (1:1000, #B2261, Sigma).

### Microtubule depolymerization

For microtubule depolymerization by cold treatment, prior to freeze-cracking and fixation, slides were incubated on metal blocks placed on ice for 20 minutes. For nocodazole treatment, L4-stage *C. elegans* worms were subjected to *perm-1(RNAi)* for 20 h at 20 °C prior to sample preparation. Worms were dissected on poly-L-lysine–coated slides in either M9 + 1% DMSO (control) or M9 + 1% DMSO supplemented with nocodazole (10 µg/ml). Following dissection, embryos were incubated in the respective buffer for 3 minutes, after which a glass coverslip was gently placed over the sample and the slide was immediately frozen for freeze cracking on dry ice. Subsequent steps of the immunofluorescence protocol were performed as described above under Indirect immunofluorescence.

### Microscopy and live-cell imaging

Indirect immunofluorescence and live-cell imaging were performed on a Leica DMi8 Stellaris 5 microscope controlled by LASX software, using a 63x HC PL APO objective (NA 1.3; glycerol immersion; WD 0.3 mm; CS2) or 100x HC PL APO objective (NA 1.4; OIL; WD 0.13 mm; CS2). Cell culture immunofluorescence Z-stack images were acquired using a Leica TCS SP5 confocal microscope with a Plan Apo 63×/1.4 NA oil-immersion objective and controlled through Leica Application Suite Advanced Fluorescence (LAS AF) software. For indirect immunofluorescence, samples were imaged at 1024×1024-pixel resolution using 0.7µm optical slices. For live-cell imaging, gravid adults were dissected to release embryos which were mounted on a 4% agar pad. Z stacks were acquired every 30 seconds laser with a z-step size of 0.71 µm. For imaging adult worms, worms were immobilized with 2.5 mM levamisole in M9 buffer and imaged using differential interference contrast (DIC) microscopy combined with fluorescence imaging. All images were analyzed using Fiji (Schindelin et al., 2012; Tinevez et al., 2017).

### Measurements of SAS-1 centriolar and satellite-like foci signal intensity, area and number

SAS-1 centriolar and satellite-like foci signa intensity measurements were performed in Fiji (Schindelin et al., 2012; Tinevez et al., 2017) after combining 2 z-stacks spanning the full centrosomal SAS-1 signal. For each centrosome, centriolar SAS-1 intensity was quantified using a fixed ROI (0.851 µm²) placed over the centriole; this region was then masked to eliminate centriolar contribution prior to satellite measurements. Satellite-associated SAS-1 intensity was subsequently measured within a larger fixed ROI (15.346 µm²) centered on the centrosome. Background signal was measured using the same ROI size in a nearby region lacking SAS-1 signal and subtracted from the corresponding intensity measurements. For satellite segmentation, images were converted to 8-bit, processed with a 2.0-pixel top-hat filter for background signal subtraction, followed by a 2-pixel Gaussian-weighted median filter for smoothing the satellite signal, and thresholded using Fiji’s default method with the lower threshold adjusted within 50–255 to outline satellite puncta. Satellite number and area were extracted using Analyze Particles, and for each centrosome the total satellite count and mean particle area were recorded, with each data point representing one centrosomal replicate. Statistical analysis was performed by testing for normality using the D’Agostino & Pearson test. Because the 2-cell and 4-cell data points come from the same embryo, a paired two-tailed t-test was used to evaluate significance between conditions.

### Colocalization of SAS-1 with SSNA-1

Colocalization between SSNA-1 and SAS-1 was analyzed using the Fiji colocalization plugin JaCoP (Bolte and Cordelieres, 2006). Images from both channels were used for analysis. For centrosomal colocalization, ROIs were defined such that the centrosome was positioned at one extreme of the ROI rather than at the center. To exclude random colocalization, one channel was rotated by 90°. Pearson’s correlation coefficient was calculated and used as a quantitative measure of colocalization. Differences between experimental and randomized conditions were evaluated using Wilcoxon signed-rank test with Holm correction.

### Venn diagram for colocalization and subset between SSNA-1 and SAS-1

To quantify colocalized and non-colocalized puncta between SSNA-1 and SAS-1, three consecutive z-slices were merged for analysis. A circular ROI (160px × 160px; corresponding to 7.22µm × 7.22µm at 0.0451µm/px) was drawn over the centrosomal region, and the ROI was split into single channels. Each channel was thresholded using Otsu with fixed intensity ranges (SSNA-1: 50–255; SAS-1: 60–255). Where puncta appeared merged, watershed separation was applied prior to particle analysis. Particle analysis was then performed to extract centroid coordinates (x, y) for all SSNA-1 and SAS-1 puncta within the ROI.

Puncta colocalization was determined by centroid-to-centroid proximity using a one-to-one (“UniqueMatch”) nearest-neighbor pairing approach. For each SSNA-1 punctum, the nearest SAS-1 punctum was identified and considered a match if the distance was ≤1 pixel (0.0451 µm). The number of accepted pairs was recorded as the UniqueMatch colocalized puncta count. From these counts, SSNA-1-only, SAS-1-only, and the shared SSNA-1/SAS-1 subset were calculated per sample. Values from n=10 total foci were summed accordingly and visualized as a Venn diagram generated in R.

### Concentric circles intensity measurements

To quantify the distribution of satellite-like structures relative to SPD-5, the Concentric Circles plug-in in Fiji was used. A total of 25 circles were generated for each measurement. The inner most circle was centered on the centriolar position and circles were drawn with a line width of 2.0 pixels, with inner and outer radii of 5 and 50 pixels, respectively. Intensity measurements were performed on Z-stacks combined into two representative projections. Background subtraction was performed by using the mean intensity of the outermost three circles, which served as the background reference. Background-corrected total intensities were calculated for each region by subtracting this value from the corresponding centrosomal and SAS-1 fluorescence intensities. A total of n=10 regions representing different cell-cycle stages were quantified, averaged and plotted using GraphPad Prism.

### Area and density measurement of *szy-20(bs52)*

PCM Mass was determined by measuring RFP::SPD-5 intensity on raw images by analyzing z-stacks using Manual tracking with the TrackMate plugin in Fiji (Schindelin et al., 2012; Tinevez et al., 2017). Metaphase was determined by the DNA condensation visualized by the mCherry::H2B marker. Fluorescent signal was measured in a sphere with a fixed radius of 2.496 µm. A sphere with the same radius was used to measure the cytoplasmic background signal and the background signal outside the embryo. For each datapoint, the cytoplasmic background signal was subtracted from the fluorescent signal. PCM area was measured at metaphase by creating a maximum *z*-projection of the RFP::SPD-5 channel and extracting the PCM by removing outliers (radius 2; threshold: 5) and despeckling the whole stack. PCM shape was then converted into black/white outlines using the ‘Otsu white’ threshold. Area was measured by the Fiji software (Schindelin et al., 2012). PCM volume was calculated from the PCM area, and density was determined by calculated PCM mass per PCM volume.

### 1,6-Hexanediol treatment

On the day of imaging, a custom imaging chamber was constructed with poly-L-lysine coated 24×60mm cover slip and double-sided tape that is punched in to create small imaging chambers (Lebedev et al., 2023). Adult *C. elegans* that were subjected to *perm-1(RNAi)* were dissected in 30 µl M9 containing Hoechst 33342 (1:1000) to assess eggshell permeabilization. Embryos were immediately transferred into the imaging chamber. 1,6-hexanediol was slowly added to the imaging chamber to a final concentration of 10%. Near the end of imaging, embryos were briefly illuminated with a 405-nm laser to control for eggshell permeabilization using Hoechst 33342. Only permeabilized embryos were used for analysis. For 1,6-hexanediol treatment of cultured cells, a 10% 1,6-hexanediol solution was prepared fresh in DMEM immediately before use. Transfected cells that were already seeded in 6-well plates containing 2 mL of culture medium were treated by removing 1 mL of medium from each well and replacing it with 1 mL of 10% 1,6-hexanediol in DMEM, resulting in a final concentration of 5% 1,6-hexanediol. Following incubation for 5 min at room temperature, cells were immediately fixed with methanol and processed for immunofluorescence as described above.

### Quantification of subcellular localization phenotypes

Cells that were transfected by different SAS-1::EGFP constructs were scored manually from fluorescence images according to the predominant localization pattern of the expressed construct. Cells exhibiting diffused cytoplasmic fluorescence were classified as cytosolic, cells with filamentous signal overlapping the microtubule network were classified as microtubule-associated, and cells containing enriched foci or aggregate-like structures were classified as accumulated. Cells were classified as cytosolic when the fluorescent signal was predominantly diffuse and fewer than five discrete foci or aggregate-like structures were observed. For each construct and treatment condition, cells were assigned to a single category by visual inspection, and the distribution of localization phenotypes was reported as a percentage of the total transfected cells counted.

### Fluorescence Recovery After Photobleaching

Embryos were mounted for live imaging as described above. FRAP was performed on a Leica Stellaris confocal microscope using a 100x objective at room temperature. Two-cell stage embryos were imaged during late anaphase to telophase, when the two centrosomes were clearly separated. A rectangular ROI (84.73 µm²; 9.48 µm × 8.94 µm) encompassing one centrosome was selected and photobleached using the 488-nm laser (80% intensity) for 15 s. The unbleached centrosome at the other spindle pole served as control. Fluorescence recovery was monitored for 30 timepoints with 22 second intervals, using identical acquisition settings. For quantification, fluorescence intensity of the pre-bleach, post-bleach, and control centrosomes was measured by integrating signal across 10 z-stacks, ensuring that all potential centrosomal satellite signal in the vicinity was captured. Centriolar SAS-1 was first measured using a small circular ROI (0.854 µm²); the centriole signal was then excluded from the image, and satellite-associated signal was measured using a larger circular ROI (15.296 µm²). Raw intensities were normalized to the pre-bleach fluorescence level and plotted to compare signal dynamics between control and bleached regions across cell-cycle stages. Signal intensity curve represents mean ± SEM values measured from n=6 embryos. The half-recovery level was defined as I_1/2_ = I_0_ +(0.5(I_∞_ + I_0_)), and t_1/2_ was taken as the first timepoint at which the normalized recovery reached or exceeded I_1/2_. The half-recovery intensity was calculated by taking the intensity values of the first time point right after FRAP as I_0_. The plateau intensity (I_∞_) was estimated as the mean of three consecutive time points within the plateau phase of the recovery curve.

### Immunoblotting

Worm protein lysates were separated by 10% SDS-PAGE and analyzed by western blotting. N- and C- terminally GFP-tagged SAS-1 proteins were probed using mouse anti-GFP antibody (mouse monoclonal 1:5000, Roche). An anti-α-tubulin antibody (mouse monoclonal 1:1000, DM1α, Sigma) was used as a loading control. Proteins of interest were detected using HRP-conjugated secondary antibodies against mouse (1:7500, Bio-Rad Laboratories). The proteins were visualized using Amersham ECL Prime Western Blotting Detection Reagent (GE Healthcare). The protein ladder used was from Proteintech (PL00001).

### Embryonic survival assay

L4 worms were singled and incubated at 15°C and 25 °C for 16-20 hours. Embryos from 60 worms were scored for each genotype (Erpf et al., 2019; Schreiner et al., 2025). The laid eggs were counted and compared to the hatched larvae after 48 hours to assess the embryonic lethality.

### Cell culture and DNA Transfection

HeLa Flp-In T-REx cells, previously described in Schneid et al., (2021), were kindly provided by the laboratory of Prof. E. Zanin and used for transient transfection and immunofluorescence staining experiments. Cells were maintained in complete DMEM (Sigma-Aldrich, #D0810-500ML) supplemented with 10% fetal bovine serum (FBS) (Bio&Sell, # FBS.HP.0500) and 1% penicillin–streptomycin (Sigma, #P4333) at 37°C and 5% CO₂. Cells were passaged every 3-4 days at approximately 70-80% confluency. For transfection, cells were detached using trypsin (Thermo Fisher, #15400054), counted, and seeded at the required density for each experiment. After 24h, the culture medium was replaced with antibiotic-free DMEM to minimize transfection-associated cytotoxicity. Cells were transfected using jetOPTIMUS® transfection reagent (Polyplus, #101000051) according to the manufacturer’s protocol. Briefly, for each well, 1 µg DNA was diluted in 200 µL jetOPTIMUS buffer, followed by the addition of 1 µL jetOPTIMUS reagent, corresponding to a 1:1 ratio of DNA (µg) to reagent (µL). The transfection mixture was incubated for 10 min at room temperature and then added dropwise to the cells. Cells were incubated with the transfection mixture overnight before subsequent experimental procedures.

### Statistical analysis

Statistical analysis was performed in R and Prism 8 software (GraphPad Software, La Jolla, CA).

For 2-cell and 4-cell embryo-averaged paired satellite area and number measurements, datasets passed the D’Agostino-Pearson normality test, and were compared by two-tailed paired t-test. For endogenous and overexpression satellite number and area analysis, statistical significance was assessed using a Mann-Whitney-U test. SSNA-1/SAS-1 colocalization data were analyzed using Wilcoxon signed-rank test with Holm correction for multiple comparisons. Statistical comparisons of embryonic viability between worm strains were analyzed using an unpaired Welch’s t-test. The percentage of centrosomes with SSNA-1 satellites was analyzed using a two-sided Fisher’s exact test. Statistical comparisons of PCM area and PCM density measurements were performed using either a Welch two-sample t-test (metaphase) or a Mann-Whitney-U test (anaphase). Normality of each dataset (control and *szy-20(bs52)* mutant) was assessed using the Shapiro-Wilk test. For datasets that followed a normal distribution (metaphase), equality of variances was evaluated using an F-test. As variances were unequal, statistical significance was determined using a Welch two-sample t-test. For datasets in which at least one group did not follow a normal distribution (anaphase), statistical significance was assessed using a Mann-Whitney-U test.

Significance levels are defined as follows: ****P < 0.0001, ***P < 0.001, **P < 0.01, *P < 0.05, NS, not significant.

## DATA AVAILABILITY

not applicable

## ACKNOWLEDGMENTS

We thank the CALM facility for support during live-cell imaging. The Leica Stellaris 5 confocal microscopes used in this study were funded by the Deutsche Forschungsgemeinschaft (DFG, German Research Foundation) – project numbers 495215303 and INST 86/2231-1 FUGG to LMU biology. We like to express our gratitude to Kevin O’Connell for sharing the OC1018 strain prior to publication, Pierre Gönczy for providing the GZ1966 strain and the codon optimized SAS-1 cDNA. Some strains were provided by the CGC, which is funded by NIH Office of Research Infrastructure Programs (P40 OD010440). We would also like to thank Özgün Bahar and Disha Shenai for building tools and conducting preliminary experiments.

## AUTHORS CONTRIBUTIONS

A. B. Tiryakiler conceptualization, data curation, formal analysis, investigation, methodology, supervision, validation, visualization, writing original draft; S. Z. A. Talib conceptualization, data curation, formal analysis, investigation, methodology, supervision, validation, visualization; A. F. Soares, F. T. Altug and A. Heim formal investigation, validation; E. Zanin conceptualization, writing original draft; T. Mikeladze-Dvali conceptualization, data curation, formal analysis, funding acquisition, investigation, methodology, project administration, resources, supervision, validation, visualization, writing origin

## VIDEOS

Video 1: SAS-1 satellite-like foci appear at the transition from two- to four-cell embryos

Video 2: FRAP of a spindle pole of a 2-cell embryo

Video 3: Embryo treated with buffer without 10% 1,6-Hexanediol

Video 4: Embryo treated with buffer with 10% 1,6-Hexanediol

Video 5: SAS-1 satellite-like foci appear in one-cell embryos when SAS-1::GFP is overexpressed

