## Supplemental Figures and Table for "The N-terminus of SAS-1 promotes microtubule-dependent centriolar satellite formation"

**SUPPLEMENTAL INFORMATION:**

**TABLES**

Supplemental Table 1: Worm strains used in this study

**Figure S1. All endogenously tagged SAS-1 transgenes localize to satellite-like foci.**

A) Schematics of endogenously tagged SAS-1 strains. B) Immunostaining of SAS-1::mkate2-expressing 2-cell embryo. C) Stills of a 1-cell embryo expressing SAS-1::GFP at metaphase, from Video 1. Same embryo as in Figure 1A. D) SAS-1::mkate2 localized to foci in multicellular embryos, amphid cilia and to cilia in the male tail. E) Embryonic survival rates of control (N2) and endogenously tagged SAS-1 strains. Significance was assessed using unpaired Welch's t-test, error bars are SD n = number of embryos and larvae analyzed. F) Analysis pipeline for quantification of satellite number and area.

Scale bars 10 $\mu$m and 2 $\mu$m in inset.

**Figure S2. SAS-1 satellite-like structures partially follow the PCM dynamics during the cell cycle.**

A) and C) Representative images of control and *szy-20(bs52)* embryos in metaphase and anaphase, at the transition from 2- to 4-cell, stained for SAS-1 and SPD-5. Red dashed line represents the outline of the PCM, magenta dashed circles indicate satellites. B) and D) Graphs showing how SAS-1 signal is distributed in respect to the PCM control and *szy-20(bs52)* embryos. E) Schematics of PCM area and density measurements in P1 cells. F) Representative images of centrosomes and their masks in metaphase and anaphase of control and *szy-20(bs52)* P1 cells. G) and H) PCM area and density measurements of P1 centrosomes in control and *szy-20(bs52)* embryos. Statistical significance was assessed using the Welch's two-sample t-test for metaphase and the Mann-Whitney-U test for anaphase, error bars are SEM.

Scale bars 10$\mu$m, n = number of centrosomes analyzed.

**Figure S3. The dynamics of SAS-1 satellite-like structures is influenced by microtubules.**

A) and B) Representative images of 4-cell embryos stained with $\alpha$-Tubulin and SAS-1, treated with DMSO (A) or nocodazole (B). C) and D) Representative images of embryos stained with $\alpha$-Tubulin and SAS-1, before (C) and after cold treatment (D). E) and F) Representative images of embryos stained for SPD-5 and SAS-1 without (E) and without (F) cold treatment. Bottom: graphs showing SAS-1 signal distribution in respect to the PCM with and without cold treatment.

Scale bars 10$\mu$m, error bars are SEM, n = number of centrosomes analyzed.

**Figure S4. SAS-1 satellite-like structures form in a dose-deepened manner.**

A) Stills of a four-cell embryo expressing endogenously tagged SAS-1::GFP and pmai-1:SAS-1::GFP at metaphase. C) Mean centriolar satellite area at centrosomes in four-cell embryos. D) Mean centriolar satellite number at centrosomes in four-cell embryos. Scale bars 10$\mu$m, significance was assessed using Mann-Whitney-U test for satellite number and area analysis, error bars are SEM, n = number of centrosomes analyzed.

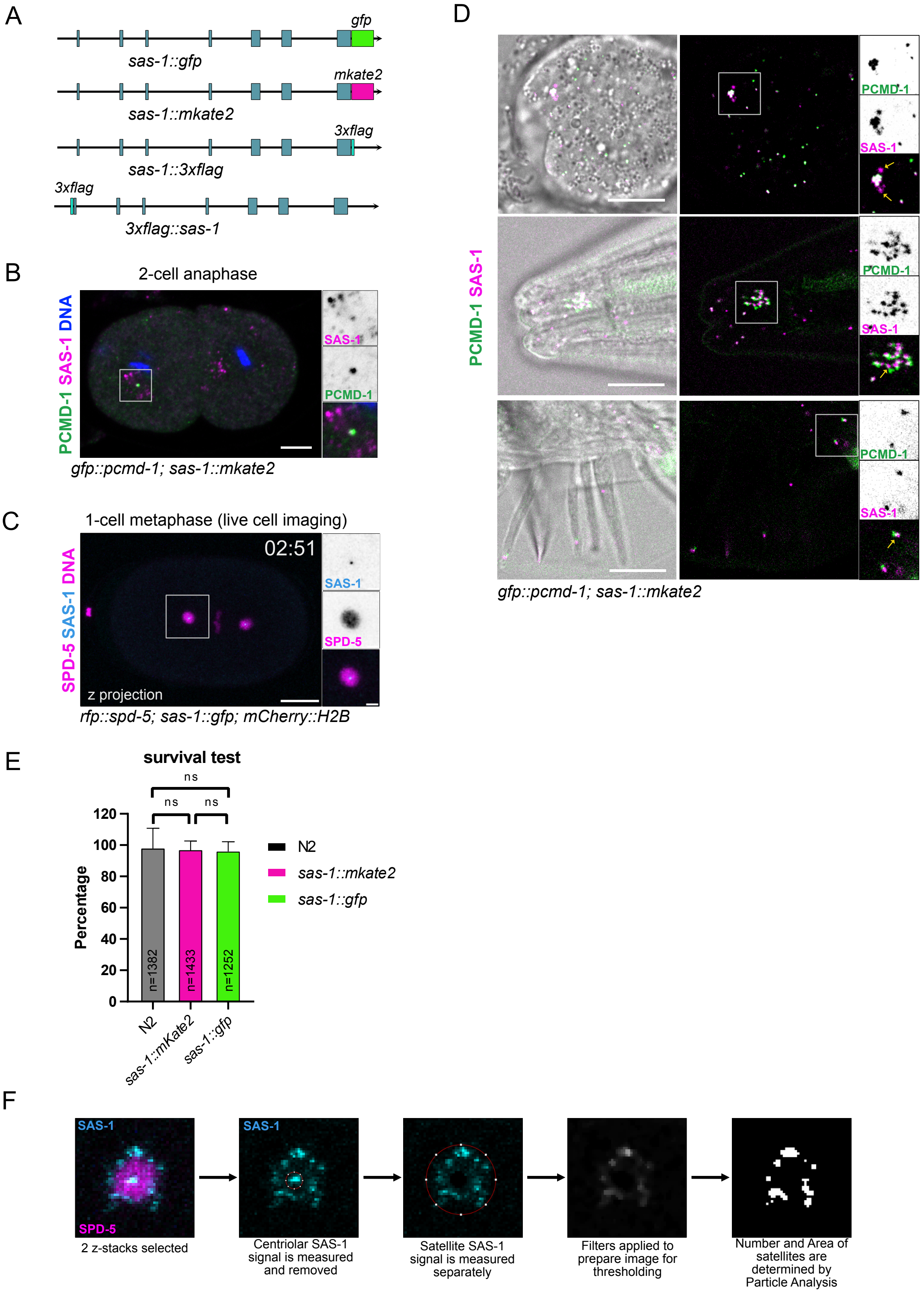
A
gfp
sas-1::gfp
mkate2
sas-1::mkate2
3xflag
sas-1::3xflag
3xflag
3xflag::sas-1
B
2-cell anaphase
PCMD-1 SAS-1 DNA
SAS-1
PCMD-1
gfp::pcmd-1; sas-1::mkate2
C
1-cell metaphase (live cell imaging)
02:51
SPD-5 SAS-1 DNA
SAS-1
SPD-5
z projection
rfp::spd-5; sas-1::gfp; mCherry::H2B
D
PCMD-1 SAS-1
PCMD-1
SAS-1
gfp::pcmd-1; sas-1::mkate2
E
survival test
ns
Percentage
N2
sas-1::mkate2
sas-1::gfp
n=1382
n=1433
n=1252
sas-1::mKate2
F
SAS-1
SPD-5
2 z-stacks selected
Centriolar SAS-1 signal is measured and removed
Satellite SAS-1 signal is measured separately
Filters applied to prepare image for thresholding
Number and Area of satellites are determined by Particle Analysis

**Figure S1**

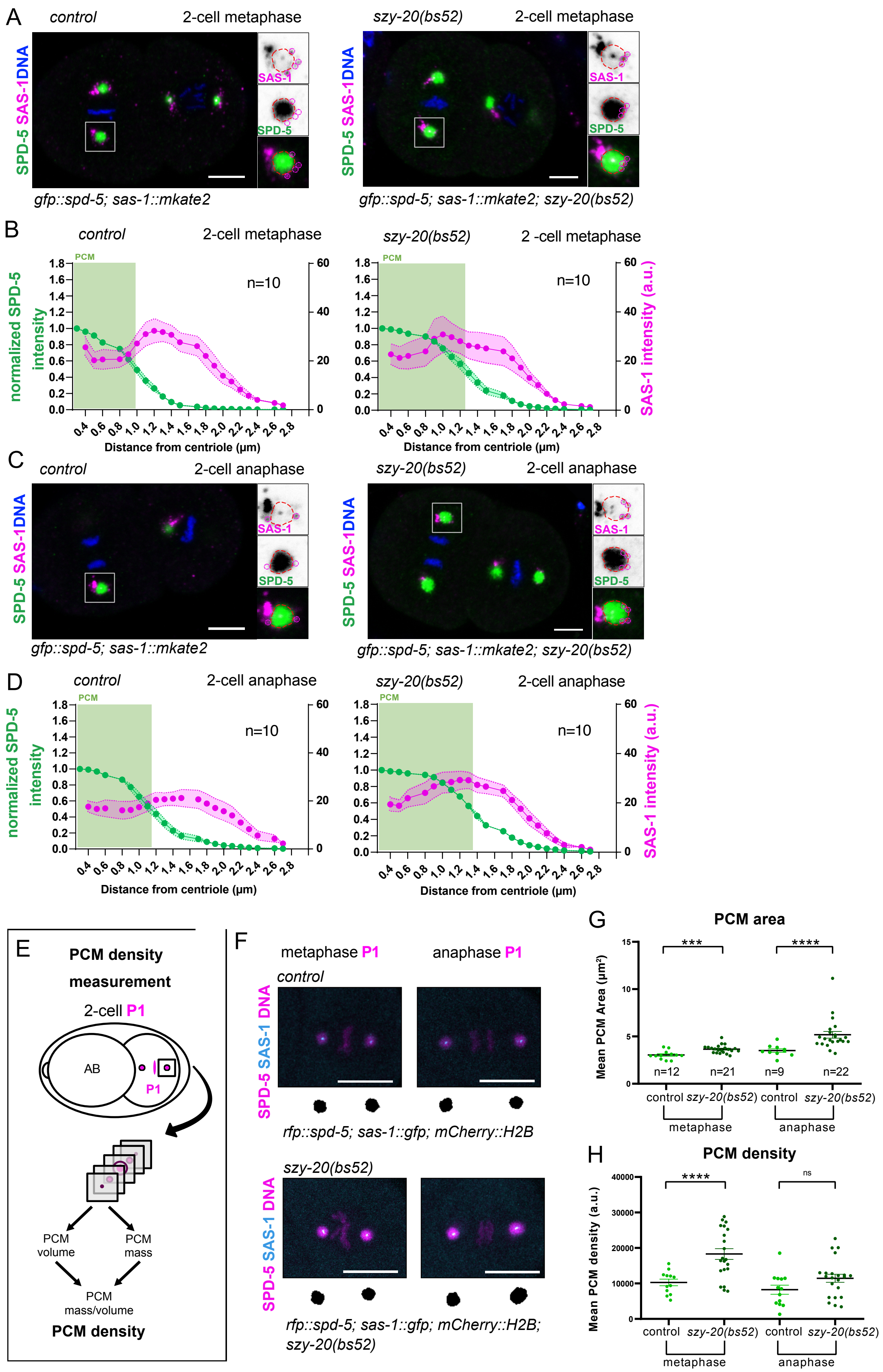
A
control
2-cell metaphase
szy-20(bs52)
2-cell metaphase
SPD-5 SAS-1 DNA
SAS-1
SPD-5
gfp::spd-5; sas-1::mkate2
gfp::spd-5; sas-1::mkate2; szy-20(bs52)
B
control
2-cell metaphase
szy-20(bs52)
2 -cell metaphase
PCM
n=10
normalized SPD-5 intensity
SAS-1 intensity (a.u.)
Distance from centriole (µm)
C
control
2-cell anaphase
szy-20(bs52)
2-cell anaphase
D
control
2-cell anaphase
szy-20(bs52)
2-cell anaphase
E
PCM density measurement
2-cell P1
AB
P1
PCM volume
PCM mass
PCM mass/volume
PCM density
F
metaphase P1
anaphase P1
control
SPD-5 SAS-1 DNA
rfp::spd-5; sas-1::gfp; mCherry::H2B
szy-20(bs52)
rfp::spd-5; sas-1::gfp; mCherry::H2B; szy-20(bs52)
G
PCM area
Mean PCM Area (µm²)
***
****
n=12
n=21
n=9
n=22
control
szy-20(bs52)
metaphase
anaphase
H
PCM density
PCM density (a.u.)
****
ns

**Figure S2**

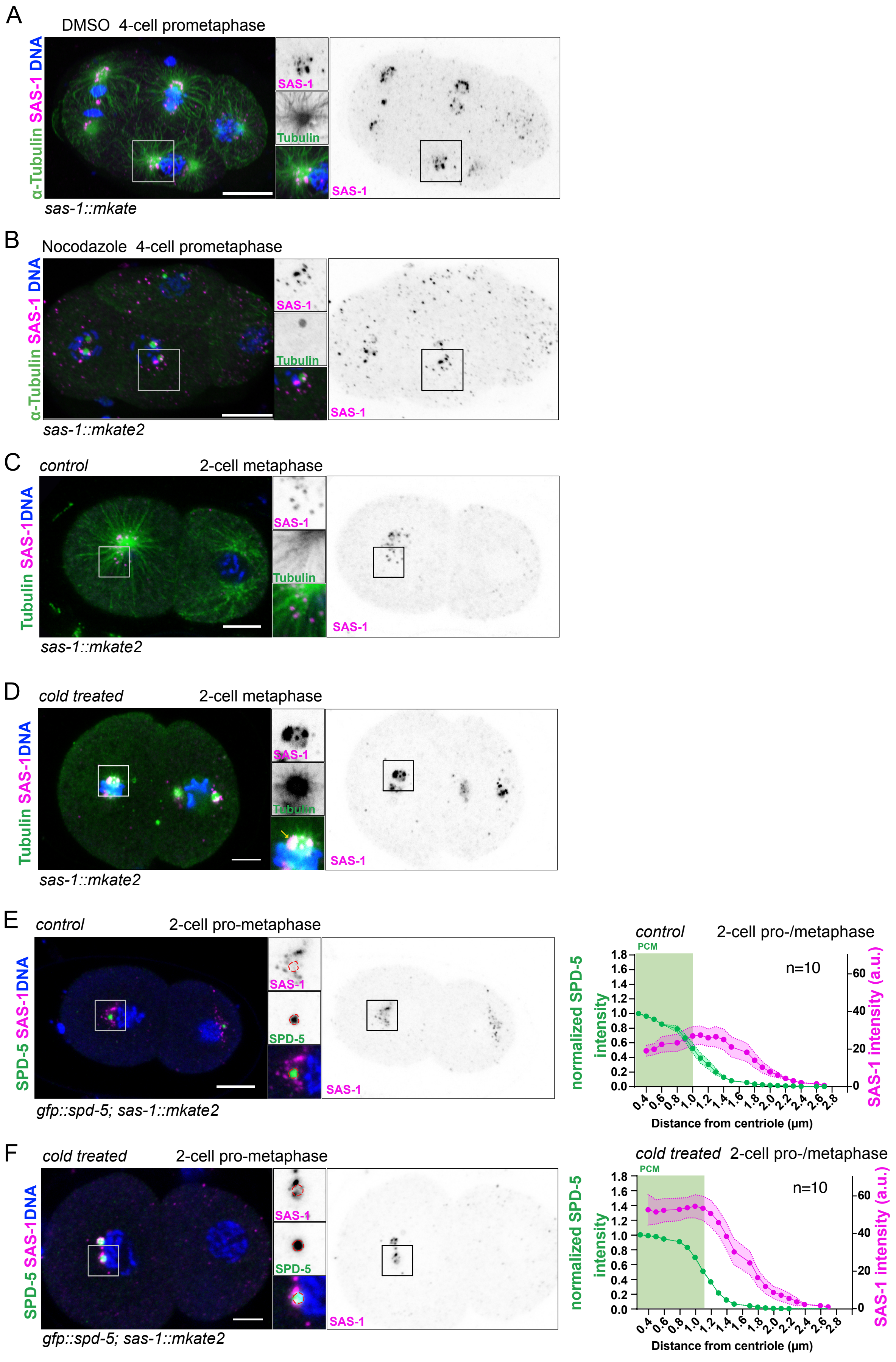

A
DMSO 4-cell prometaphase
α-Tubulin SAS-1 DNA
SAS-1
Tubulin
SAS-1
sas-1::mkate
B
Nocodazole 4-cell prometaphase
α-Tubulin SAS-1 DNA
SAS-1
Tubulin
SAS-1
sas-1::mkate2
C
control
2-cell metaphase
Tubulin SAS-1DNA
SAS-1
Tubulin
SAS-1
sas-1::mkate2
D
cold treated
2-cell metaphase
Tubulin SAS-1DNA
SAS-1
Tubulin
SAS-1
sas-1::mkate2
E
control
2-cell pro-metaphase
SPD-5 SAS-1DNA
SAS-1
SPD-5
SAS-1
gfp::spd-5; sas-1::mkate2
control
2-cell pro-/metaphase
PCM
n=10
normalized SPD-5 intensity
SAS-1 intensity (a.u.)
Distance from centriole (µm)
F
cold treated
2-cell pro-metaphase
SPD-5 SAS-1DNA
SAS-1
SPD-5
SAS-1
gfp::spd-5; sas-1::mkate2
cold treated
2-cell pro-/metaphase
PCM
n=10
normalized SPD-5 intensity
SAS-1 intensity (a.u.)
Distance from centriole (µm)


**Figure S3**

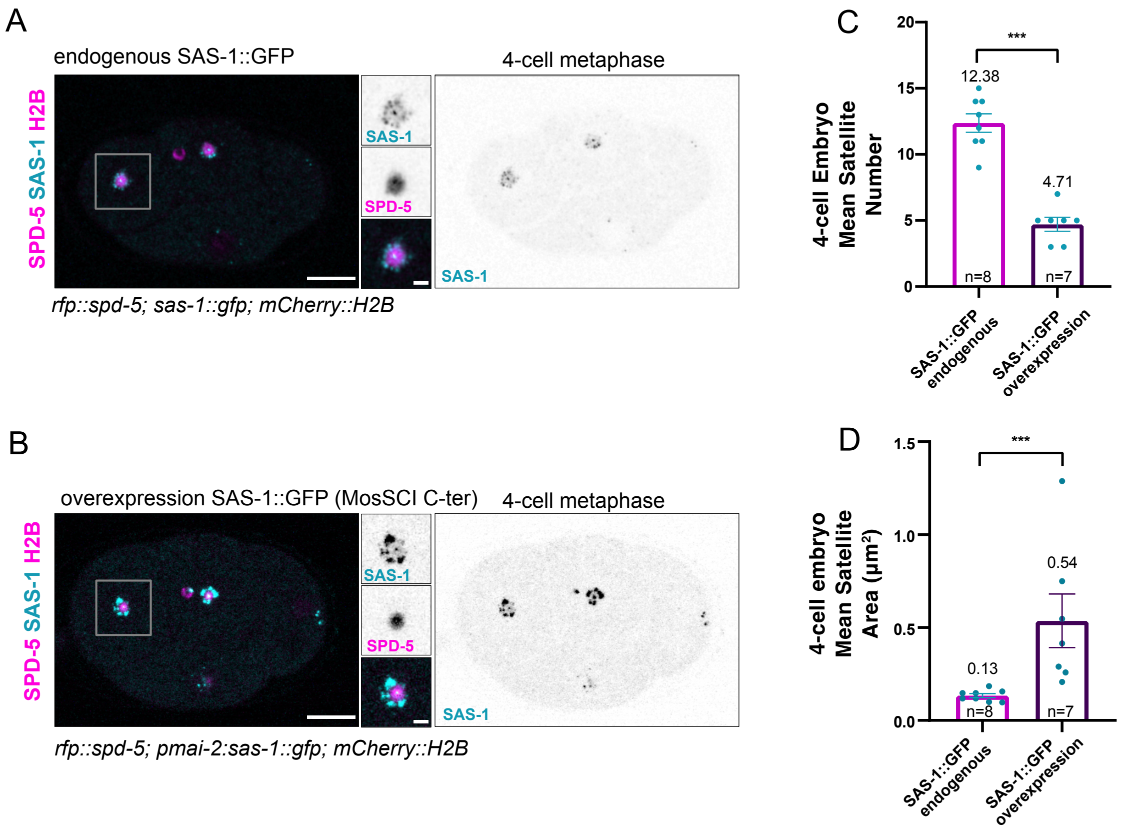
A
endogenous SAS-1::GFP
4-cell metaphase
SPD-5 SAS-1 H2B
SAS-1
SPD-5
SAS-1
rfp::spd-5; sas-1::gfp; mCherry::H2B
B
overexpression SAS-1::GFP (MosSCI C-ter)
4-cell metaphase
SPD-5 SAS-1 H2B
SAS-1
SPD-5
SAS-1
rfp::spd-5; pmai-2:sas-1::gfp; mCherry::H2B
C
4-cell Embryo Mean Satellite Number
***
12.38
4.71
n=8
n=7
SAS-1::GFP endogenous
SAS-1::GFP overexpression
D
4-cell embryo Mean Satellite Area (µm²)
***
0.54
0.13
n=8
n=7
SAS-1::GFP endogenous
SAS-1::GFP overexpression

**Figure S4**

### Supplemental Table 1: Worm Strains

| Strain | Genotype | Source |
|---|---|---|
| GZ1966 | *sas-1(is6[sas-1::3xflag]) III* | Woglar A. et al. 2022 |
| GZ1934 | *sas-1(is7[3xflag::sas-1]) III* | Woglar A. et al. 2022 |
| OC1018 | *ssna-1(bs206 [ssna-1::spot]) IV* | Pfister J. A. et al. 2025 |
| TMD194 | *pcmd-1(syb486[gfp::pcmd-1])I; sas-1(syb2290[sas-1::mkate2]) III* | This study |
| TMD231 | *ltSi202[pVV103/pOD1021; Pspd-2::GFP::SPD-5 RNAi-resistant; cb-unc-119(+)]II; sas-1(syb2290[sas-1::mkate2]) III* | This study |
| TMD278 | *sas-1(syb2290[sas-1::mkate2]) III ruIs32 [pie-1p::GFP::H2B + unc-119(+)] III; ojIs1 [pie-1p::GFP::tbb-2 + unc-119(+)]* | This study |
| TMD388 | *szy-20(bs52) ltSi202 [pVV103; Pspd-2::GFP::SPD-5 reencoded; cb-unc-119(+)]II; sas-1(syb2290[sas-1::mkate2]) III* | This study |
| TMD373 | *sas-1(syb2290[sas-1::mkate2]) III; ssna-1(bs206 [ssna-1::spot]) IV* | This study |
| TMD397 | *spd-5(wow36[tagrfp-t^3xmyc::spd-5])I; sas-1(syb9121[sas-1::GFP])III ; ltIs37 [(pAA64) pie-1p::mCherry::his-58 + unc-119(+)] IV* | This study |
| TMD412 | *spd-5(wow36[tagrfp-t^3xmyc::spd-5])I; szy-20(bs52)II; sas-1(syb9121[sas-1::GFP])III ; ltIs37 [(pAA64) pie-1p::mCherry::his-58 + unc-119(+)] IV* | This study |
| TMD414 | *unc-32(e189) sas-1 (t1476)/hT2 III; ssna-1(bs206 [ssna-1::spot]) IV* | This study |
| TMD415 | *wt/hT2 I; sas-1(syb4965)/hT2 III; ssna-1(bs206 [ssna-1::spot]) IV* | This study |
| TMD451 | *sas-1(syb9121[sas-1::GFP])III; ocfIs2 [pie-1p:mCherry::sp12::pie-1 3'UTR + unc-119(+)]* | This study |
| TMD471 | *spd-5(wow36[tagrfp-t^3xmyc::spd-5]); mikSi61[pmai-2:sas-1(opt)::GFP] II ; ltIs37 [(pAA64) pie-1p::mCherry::his-58 + unc-119(+)] IV* | This study |
| TMD472 | *spd-5(wow36[tagrfp-t^3xmyc::spd-5]); mikSi76[pmai-2:GFP::sas-1(opt)]II; ltIs37 [(pAA64) pie-1p::mCherry::his-58 + unc-119(+)] IV* | This study |

| | | |
|---|---|---|
| TMD476 | *spd-5(wow36[tagrfp-t^3xmyc::spd-5])I/hT2 I; mikSi61[pmai-2:sas-1(opt)::GFP] II ; sas-1(syb4965)/hT2 III; ltIs37 [(pAA64) pie-1p::mCherry::his-58 + unc-119(+)] IV* | This study |
| TMD477 | *spd-5(wow36[tagrfp-t^3xmyc::spd-5])I/hT2 I; mikSi76[pmai-2:GFP::sas-1(opt)]II; sas-1(syb4965)/hT2 III; ltIs37 [(pAA64) pie-1p::mCherry::his-58 + unc-119(+)] IV* | This study |
